# ATF7IP2/MCAF2 directs H3K9 methylation and meiotic gene regulation in the male germline

**DOI:** 10.1101/2023.09.30.560314

**Authors:** Kris G. Alavattam, Jasmine M. Esparza, Mengwen Hu, Ryuki Shimada, Anna R. Kohrs, Hironori Abe, Yasuhisa Munakata, Kai Otsuka, Saori Yoshimura, Yuka Kitamura, Yu-Han Yeh, Yueh-Chiang Hu, Jihye Kim, Paul R. Andreassen, Kei-ichiro Ishiguro, Satoshi H. Namekawa

**Affiliations:** Reproductive Sciences Center, Division of Developmental Biology, Cincinnati Children’s Hospital Medical Center, Cincinnati, Ohio 45229, USA; Basic Sciences Division, Fred Hutchinson Cancer Center, Seattle, Washington 98109, USA; Department of Microbiology and Molecular Genetics, University of California, Davis, California 95616, USA; Department of Chromosome Biology, Institute of Molecular Embryology and Genetics (IMEG), Kumamoto University, Kumamoto, 860-0811, Japan; Laboratory of Chromosome Dynamics, Institute of Molecular and Cellular Biosciences, University of Tokyo, 1-1-1, Yayoi, Tokyo, 113-0032, Japan; Division of Experimental Hematology and Cancer Biology, Cincinnati Children’s Hospital Medical Center, Cincinnati, Ohio 45229, USA; Department of Pediatrics, University of Cincinnati College of Medicine, Cincinnati, Ohio 49229, USA

**Author notes:** Corresponding authors: Kei-ichiro Ishiguro, Satoshi H. Namekawa. These authors contributed equally to this work (Kris G. Alavattam, Jasmine M. Esparza, Mengwen Hu, Ryuki Shimada).

**Keywords:** Meiosis, Constitutive Heterochromatin, H3K9me3, Meiotic Sex Chromosome Inactivation, Gene activation, ATF7IP2/MCAF2

## Abstract

H3K9 tri-methylation (H3K9me3) plays emerging roles in gene regulation, beyond its accumulation on pericentric constitutive heterochromatin. It remains a mystery why and how H3K9me3 undergoes dynamic regulation in male meiosis. Here, we identify a novel, critical regulator of H3K9 methylation and spermatogenic heterochromatin organization: the germline-specific protein ATF7IP2 (MCAF2). We show that, in male meiosis, ATF7IP2 amasses on autosomal and X pericentric heterochromatin, spreads through the entirety of the sex chromosomes, and accumulates on thousands of autosomal promoters and retrotransposon loci. On the sex chromosomes, which undergo meiotic sex chromosome inactivation (MSCI), the DNA damage response pathway recruits ATF7IP2 to X pericentric heterochromatin, where it facilitates the recruitment of SETDB1, a histone methyltransferase that catalyzes H3K9me3. In the absence of ATF7IP2, male germ cells are arrested in meiotic prophase I. Analyses of ATF7IP2-deficient meiosis reveal the protein’s essential roles in the maintenance of MSCI, suppression of retrotransposons, and global upregulation of autosomal genes. We propose that ATF7IP2 is a downstream effector of the DDR pathway in meiosis that coordinates the organization of heterochromatin and gene regulation through the spatial regulation of SETDB1-mediated H3K9me3 deposition.

## Introduction

Constitutive heterochromatin forms mainly at pericentromeres and is maintained to ensure genome stability. A hallmark of constitutive heterochromatin is histone H3K9 tri-methylation (H3K9me3) (Saksouk et al. 2015). It was initially considered a static histone mark due to its stable accumulation on tandem satellite repeats at pericentric heterochromatin (PCH); however, a growing literature reveals that H3K9me3—particularly H3K9me3 mediated by the histone methyltransferase SETDB1—has broad, dynamic roles in suppressing developmental regulator genes and endogenous retroviruses in embryonic stem cells (Bilodeau et al. 2009; Matsui et al. 2010), thereby defining cellular identities in somatic development (Becker et al. 2016; Nicetto and Zaret 2019).

An essential factor in the germline, SETDB1 is required for gene regulation, the suppression of transposable elements (TEs), and the control of meiotic chromosome behavior (Liu et al. 2014; Hirota et al. 2018; Mochizuki et al. 2018; Cheng et al. 2021). The redundant H3K9me3 methyltransferases SUV39H1 and SUV39H2 are also required for male meiosis (Peters et al. 2001). Thus, the regulation of H3K9me3 is critical in male meiosis, where constitutive heterochromatin is remodeled to undergo synapsis and meiotic recombination on homologous chromosomes (Scherthan et al. 2014; Berrios 2017; Maezawa et al. 2018a). However, it remains a mystery why and how H3K9me3 undergoes dynamic regulation in male meiosis.

In addition to its roles at PCH, H3K9me3 is subject to dynamic temporal and spatial regulation on the male sex chromosomes as they undergo meiotic sex chromosome inactivation (MCSI) (Turner 2015; Alavattam et al. 2021). An essential event in the male germline, MSCI is initiated and maintained by a DNA damage response (DDR) pathway (Ichijima et al. 2011; Royo et al. 2013; Abe et al. 2022). Downstream of the DDR, SETDB1 establishes H3K9me3 on the sex chromosome and regulates MSCI (Hirota et al. 2018). SETDB1 is expressed in a broad range of cells, but there is a major knowledge gap as to how SETDB1 and H3K9me3 function in meiosis.

Here, we identify Activating transcription factor 7 interacting protein 2 (ATF7IP2), also known as MBD1-containing chromatin-associated factor 2 (MCAF2), as a novel, critical regulator of SETDB1’s spatiotemporal activity, H3K9 methylation, and global spermatogenic gene regulation. We identified ATF7IP2 based on its gene expression in the germline. In the midst of our investigation, an IP-mass spectrometry analysis identified ATF7IP2 as a SETDB1-binding protein (Hirota et al. 2018), lending the factor further contextual significance. In mitotically cycling cells, its homolog ATF7IP (MCAF1) regulates SETDB1 for H3K9me3 establishment and transcriptional silencing (Ichimura et al. 2005; Timms et al. 2016; Tsusaka et al. 2019; Tsusaka et al. 2020). We show that ATF7IP2 is a counterpart to ATF7IP that is highly expressed in the germline and essential in male meiosis, revealing roles for ATF7IP2 in MSCI, global meiotic gene regulation, and the fine-tuning of retrotransposon-derived loci such as endogenous retroviruses. By uncovering ATF7IP2’s germline functions, our study clarifies the regulatory logic for dynamic H3K9me3 deposition—and thus heterochromatin—in the male germline.

## Results

### ATF7IP2 is highly expressed in male meiosis and accumulates on heterochromatin

To understand the meiosis-specific regulation of H3K9me3, we focused on *Atf7ip2* (*Mcaf2*), a gene that is highly expressed in male meiosis as evidenced in RNA-seq datasets for germ cell development and spermatogenesis (Seisenberger et al. 2012; Hasegawa et al. 2015; Maezawa et al. 2018b) (Fig. 1A). *Atf7ip2* expression is low in male germ cells until the stage of meiosis, at which point it is highly upregulated in meiotic pachytene spermatocytes (Fig. 1A). On the other hand, its homolog *Atf7ip* (*Mcaf1*), which functions in mitotically dividing/somatic cells (Ichimura et al. 2005; Timms et al. 2016), is highly expressed in primordial germ cells and spermatogonia but is downregulated in pachytene spermatocytes. Among various tissues, *Atf7ip2* is highly expressed in testes (Supplemental Fig. S1A). Furthermore, mouse ATF7IP2 has high homology with human ATF7IP2 (Supplemental Fig. S1B), except for its long N-terminal amino acid tail, and ATF7IP2 is highly expressed in human testes’ meiotic spermatocytes (Supplemental Fig. S1C). These results raise the possibility that ATF7IP2 is an evolutionarily conserved counterpart to ATF7IP that is highly expressed in late stages of spermatogenesis.

**Figure 1.**
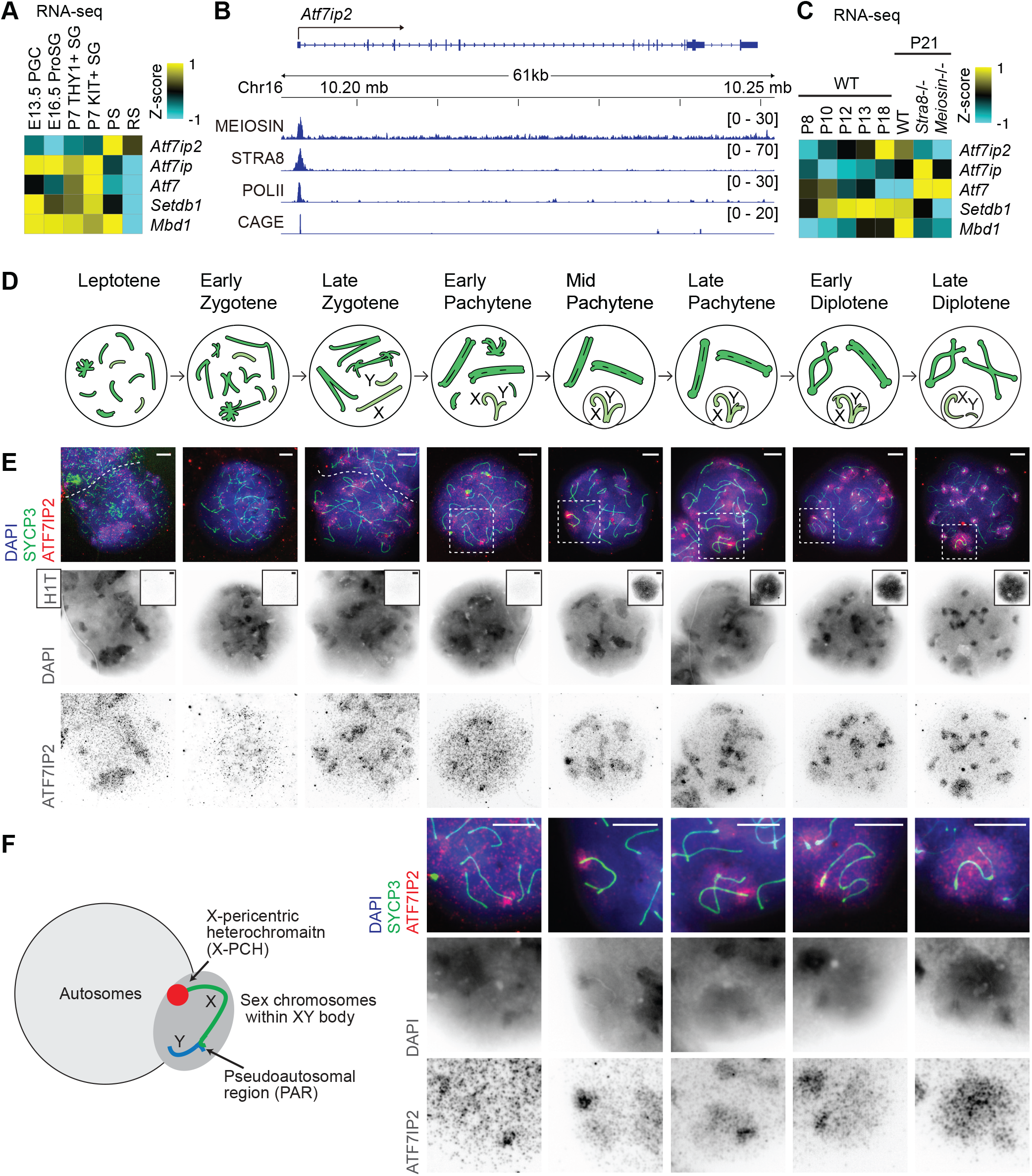
ATF7IP2 is highly expressed in male meiosis and accumulates on heterochromatin. **(A, C)** Heatmaps showing bulk RNA-seq gene expression levels across a male-germline time course for *Atf7ip2* and related genes. PGC: Primordial germ cells, ProSG: prospermatogonia, SG: spermatogonia, PS: pachytene spermatocytes, RS: round spermatids. Original data are from (Seisenberger et al. 2012; Hasegawa et al. 2015; Maezawa et al. 2018b) for (**A**) and (Ishiguro et al. 2020) for (**C**) **(B)** Track views for MEIOSIN (preleptotene-enriched testes), STRA8 (preleptotene-enriched testes), and RNA polymerase II (POLII; postnatal day (P) 10.5 testes) ChIP-seq data, and CAGE (P10.5 testes). Numbers in brackets: ranges of normalized coverage. **(D)** Schematic: chromosome behavior in meiotic prophase I of male *Mus musculus*. Darker green: autosomes; lighter green: sex chromosomes. **(E)** Meiotic chromosome spreads stained with DAPI and antibodies raised against ATF7IP2, SYCP3, and H1T; spreads represent stages of meiotic prophase I. Insets: H1T immunostaining; H1T is a nuclear marker that appears in mid pachytene nuclei and persists into haploid spermatids. SYCP3 is a marker of meiotic chromosome axes. Dashed squares are magnified in panel **F**. Scale bars: 5 μm. **(F)** Schematic: sex chromosome configuration in male meiosis. Right: magnified images of sex chromosomes from panel **E**. Scale bars: 5 μm.

To understand the regulatory mechanism for *Atf7ip2* expression, we examined the genomic distribution of MEIOSIN and STRA8, both transcription factors that heterodimerize to initiate meiosis-specific transcription (Kojima et al. 2019; Ishiguro et al. 2020). We observed MEIOSIN and STRA8 peaks at the *Atf7ip2* transcription start site (TSS) in preleptotene-enriched testes (the preleptotene stage is a liminal stage for germ cells transitioning from mitosis to meiotic prophase I) [Fig. 1B, reanalysis of (Ishiguro et al. 2020)]. These peaks coincide with the accumulation of RNA polymerase II (POLII) and Cap Analysis of Gene Expression (CAGE) signals in postnatal day 10.5 (P10.5) testes, which are enriched for preleptotene spermatocytes [Fig. 1B, reanalysis of (Li et al. 2013)]. In support of a role for MEIOSIN and STRA8 in *Atf7ip2* expression, *Atf7ip2* was downregulated in *Stra8*^-/-^ and *Meiosin*^-/-^ testes at P21 (Fig. 1C). In mouse testes, the first wave of meiosis occurs semi-synchronously, and *Atf7ip2* expression is at its highest in P18 testes, when late stages of meiotic prophase I spermatocytes first appear (Fig. 1C). Taken together, these results demonstrate that the expression of *Atf7ip2* is upregulated by MEIOSIN and STRA8, occurring amid a broad range of meiotic transcription (Kojima et al. 2019; Ishiguro et al. 2020).

To better understand the potential function of ATF7IP2, we investigated ATF7IP2 protein localization during stages of mouse male meiosis by performing immunofluorescence microscopy with chromosome spreads (Fig. 1D, E). In the leptotene stage of meiotic prophase I, when meiotic chromosome axes begin to condense, ATF7IP2 localizes on DAPI-discernible heterochromatin. ATF7IP2 continues to localize on all DAPI-discernible PCH through the zygotene stage, when homologs undergo synapsis; the pachytene stage, when homologs have completed synapsis; and the diplotene stage, when homologs begin desynapsis (Fig. 1E). Meiotic nuclei were staged through observations of chromosome axes as identified by SYCP3 (Alavattam et al. 2016; Alavattam et al. 2018), a component of meiotic axes, and the presence of the testis-specific histone variant H1T, which appears in mid pachytene nuclei and persists into haploid spermatids (Inselman et al. 2003). At the onset of the pachytene stage, the unsynapsed sex chromosomes undergo MSCI, and the most intense ATF7IP2 signals were observed on X-chromosome PCH (X-PCH) at that time (Fig. 1E, F). In the early and mid pachytene stages, ATF7IP2 localizes primarily on X-PCH; from the late pachytene stage onward, ATF7IP2 gradually spreads across the entirety of the sex chromosome domain (also referred to as the “XY domain” or “XY chromatin,”). Thus, ATF7IP2 exhibits two distinct localization patterns in meiotic prophase I: one is on the PCH of all chromosomes, and the other is intense accumulation on X-PCH that proceeds to spread through the entirety of the XY chromatin.

### ATF7IP2 is required for male meiosis

To test the function of ATF7IP2, we performed CRISPR-mediated genome editing to generate *Atf7ip2* knockout mice. We targeted a guide RNA to a site within exon 4 (Fig. 2A), which encodes a portion of the SETDB1-binding domain (SETDB1-BD) that is conserved between ATF7IP2 and ATF7IP (Fig. 2B). We obtained three alleles with deletion lengths of, respectively, 17, 31, and 169 bp. All caused *Atf7ip2* frameshift mutations, and all three homozygous *Atf7ip2* mutants displayed consistent and obvious testicular defects. For subsequent analyses, we selected the 17 bp-deletion allele as a representative; hereafter, the homozygous 17 bp-allele model is denoted *Atf7ip2*^-/-^. *Atf7ip2*^-/-^ male mice were viable but infertile, and had much smaller testes compared to littermate controls (Fig. 2C, D, E). We confirmed the depletion of ATF7IP2 proteins in *Atf7ip2*^-/-^ spermatocytes via immunofluorescence microscopy (Supplemental Fig. S2). Analyses of testicular tissue sections showed that *Atf7ip2*^-/-^ testes were devoid of haploid spermatids, and seminiferous tubules were smaller than in control testes (Fig. 2F). However, *Atf7ip2*^-/-^ spermatocytes reached the stage when H1T is enriched, the mid pachytene stage, indicating that *Atf7ip2*^-/-^ spermatocytes are arrested and eliminated in meiotic prophase I. Unlike *Atf7ip2*^-/-^ males, *Atf7ip2*^-/-^ female mice were fertile and, when crossed with *Atf7ip2*^+/-^ males, gave birth at Mendelian ratios (Supplemental Fig. S3). These results suggest that the *Atf7ip2*^-/-^ phenotype is caused by an essential, male-specific event in the germline that has gone defective.

**Figure 2.**
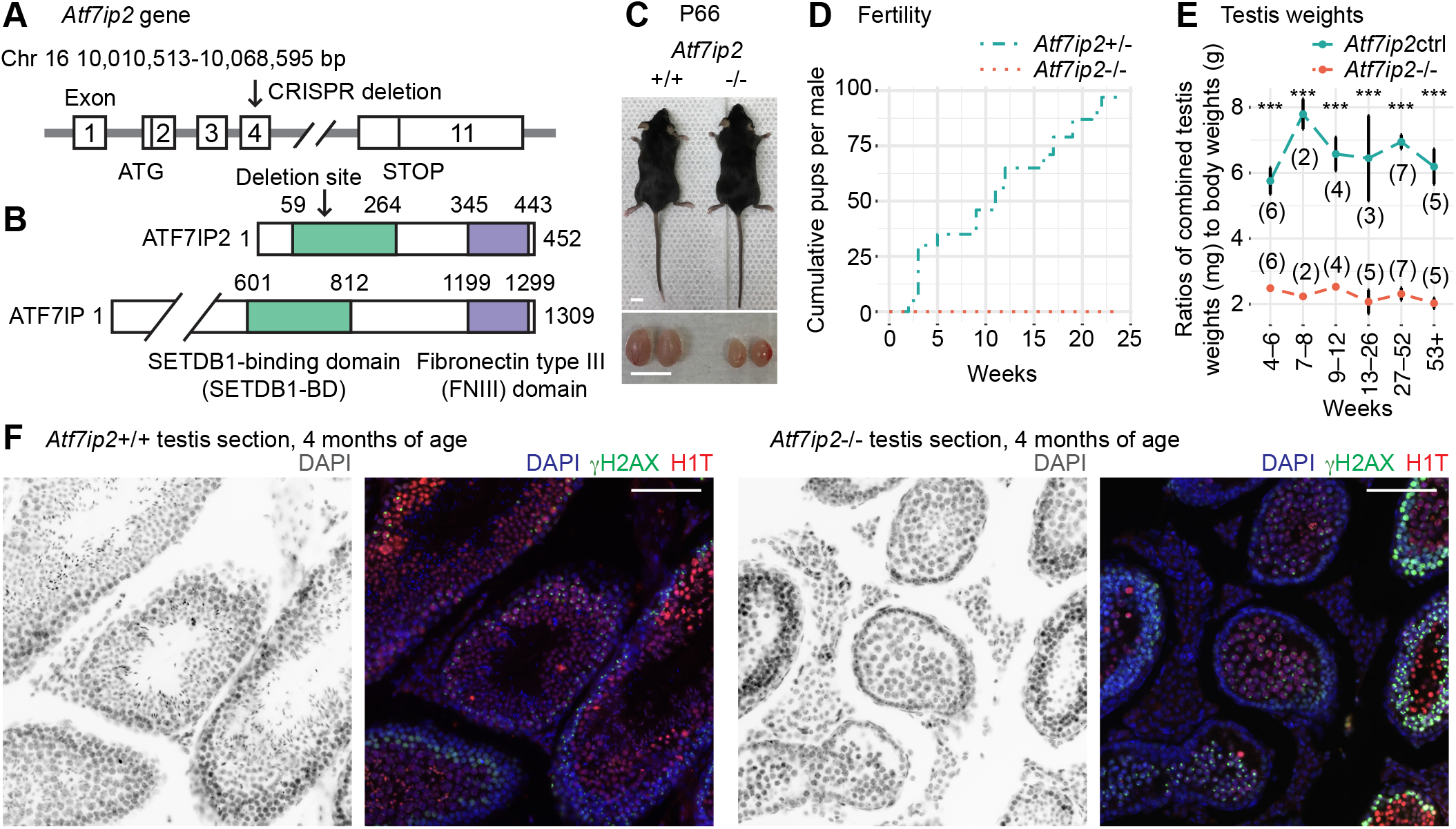
ATF7IP2 is required for male fertility. **(A)** Schematic: mouse *Atf7ip2* gene and the location of the CRISPR-mediated deletion. **(B)** Schematic: mouse ATF7IP2 and ATF7IP proteins, and their functional domains. **(C)** *Atf7ip2^+/+^* and *Atf7ip2^-/-^* males, and their testes, at postnatal day 66 (P66). Scale bars: 10 mm. **(D)** Cumulative numbers of pups sired with *Atf7ip2+/-* and *Atf7ip2-/-* males. **(E)** Testis weights for *Atf7ip2^-/-^*males and littermate controls (*Atf7ip2* ctrl: *Atf7ip2^+/+^*and *Atf7ip2^+/-^*). Numbers of independent mice analyzed are shown in parentheses. P-values are from pairwise t-tests adjusted with Benjamini-Hochberg post-hoc tests: *** < 0.001. Data are presented as mean ± SEM. **(F)** Testis sections from *Atf7ip2*^+/+^ and *Atf7ip2^-/-^* mice at 4 months of age stained with DAPI and antibodies raised against ATF7IP2, γH2AX (a marker of the DNA damage response), and H1T (a marker of germ cells in mid pachytene and subsequent stages). Scale bars: 100 μm.

### Meiotic phenotypes in male Atf7ip2^-/-^ mice

To determine the function of ATF7IP2, we characterized the meiotic phenotype of *Atf7ip2*^-/-^ male mice in detail. We performed immunostaining to analyze chromosome spreads from *Atf7ip2*^-/-^ testes for a specific marker of the DDR: phosphorylated Serine 139 of the histone variant H2AX (γH2AX). In the leptotene and zygotene stages, the DDR/checkpoint kinase Ataxia Telangiectasia Mutated (ATM) triggers the formation of γH2AX domains throughout nuclei in response to programmed double-stranded breaks (DSBs; induced by the topoisomerase-related enzyme SPO11); with the completion of DNA repair and concomitant autosomal synapsis, γH2AX disappears from autosomes (Mahadevaiah et al. 2001; Bellani et al. 2005). In the latter steps of this process, Ataxia Telangiectasia and Rad3-Related (ATR), another DDR/checkpoint kinase, mediates γH2AX formation on unsynapsed chromatin; in normal pachytene nuclei, this results in the confinement of γH2AX to the unsynapsed XY chromosomes, an essential event in the initiation of MSCI (Royo et al. 2013). Thus, γH2AX staining, together with SYCP3 staining, provides key insights into general meiotic phenotypes (Abe et al. 2018; Alavattam et al. 2018). In *Atf7ip2*^-/-^ spermatocytes, pan-nuclear γH2AX formation occurs normally in the early/mid zygotene stage (Fig. 3A), and relative populations of zygotene spermatocytes are comparable between *Atf7ip2*^-/-^ testes and littermate controls (Fig. 3B). In the *Atf7ip2*^-/-^ pachytene spermatocytes, γH2AX formation on the XY chromosomes takes place (Fig. 3A); however, we noted a significant increase in the relative population of early/mid pachytene spermatocytes, while diplotene spermatocytes were rare and largely depleted from *Atf7ip2*^-/-^ testes (Fig. 3B). These analyses suggest that ATF7IP2 has a critical function as spermatocytes transition from the pachytene to diplotene stages.

**Figure 3.**
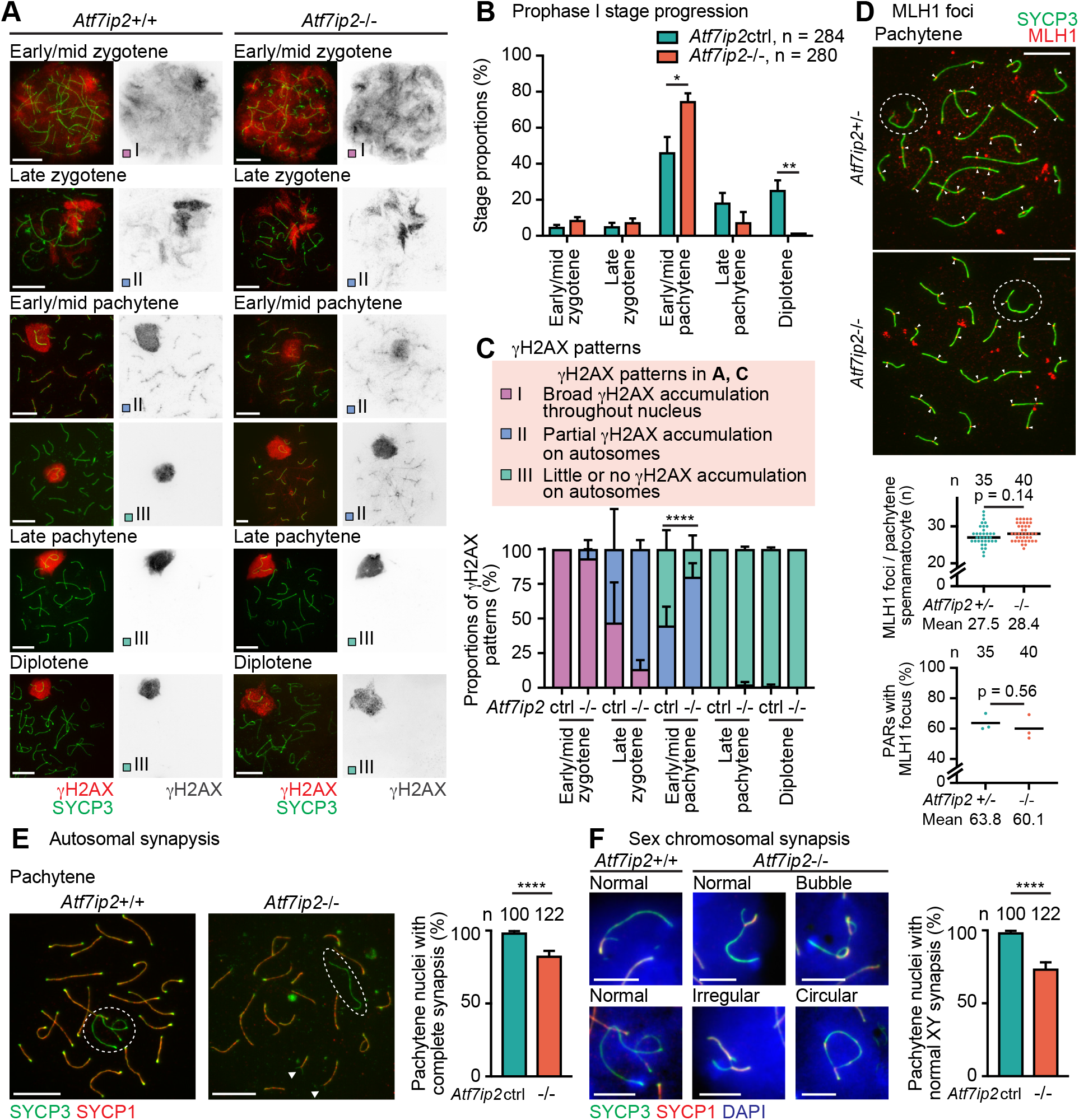
DDR and chromosome synapsis are mildly impaired in *Atf7ip2^-/-^* spermatocytes. **(A)** *Atf7ip2^+/+^* and *Atf7ip2^-/-^* spermatocyte chromosome spreads stained with antibodies raised against SYCP3 and γH2AX. γH2AX accumulation patterns are one of three classifications described in panel **C**. Scale bars: 10 μm. **(B)** Meiotic prophase I stage populations quantified as mean ± SEM for three independent littermate pairs. Numbers of analyzed nuclei are indicated. Data are from five independent littermate pairs at P44, P56, P66, P66, and P69. P-values are from unpaired two-tailed t-tests: * < 0.05, ** < 0.01. **(C)** Stage-wise proportions of γH2AX accumulation patterns for three independent littermate pairs. Patterns are classified with three criteria (see top). P-values are from Fisher’s exact tests: **** < 0.0001. **(D)** Chromosome spreads stained with antibodies raised against SYCP3 and MLH1. Arrowheads indicate MLH1 foci. Dot plot (top): distributions of MLH1 counts from three independent littermate pairs. Dot plot (bottom): proportions of MLH1 focus-associated XY pseudoautosomal regions (PARs) from three independent littermate pairs. Numbers of analyzed nuclei are indicated. Data are from three independent littermate pairs at P108, P115, P122. Bars represent means. P-values are from unpaired t-tests. **(E, F)** Chromosome spreads stained with antibodies raised against SYCP3 (a marker of all chromosome axes) and SYCP1 (a marker of only synapsed axes). Scale bars: 10 μm (**E**), 5 μm (**F**). Bar plots: proportions of pachytene nuclei with normal synapsis of autosomes (**E**) and sex chromosomes (**F**). Data are from four independent littermate pairs at P44, P66, P66, and P69, and presented as mean ± SEM. P-values are from unpaired t-tests: * < 0.05, ** < 0.01.

Following our established criteria for SYCP3- and γH2AX-based meiotic staging (Abe et al. 2018; Alavattam et al. 2018), we analyzed γH2AX staining patterns in more detail. In *Atf7ip2*^-/-^ early/mid pachytene spermatocytes, the removal of γH2AX from autosomes was delayed in comparison to controls (Fig. 3C). In normal meiosis, γH2AX accumulates through the whole of leptotene and early zygotene nuclei (Pattern I, Fig. 3A and 3C); as spermatocytes progress into the late zygotene stage, γH2AX accumulation transitions from a pan-nuclear diffuse signal to concentrated accumulation on the chromatin associated with unsynapsed chromosome axes, albeit with partial signals remaining along synapsed autosomes (Pattern II); by the mid and late pachytene stages, γH2AX is confined to XY chromatin, having largely disappeared from autosomes (Pattern III). In *Atf7ip2*^-/-^ early/mid pachytene spermatocytes, γH2AX remains on autosomes longer than in littermate controls (Fig. 3C), suggesting that, in the absence of ATF7IP2, autosomal DDR signaling is affected.

Following on this, we investigated the outcome of meiotic recombination by scoring the numbers of MLH1 foci—which illuminate crossover sites—on chromosome axes. Numbers of MLH1 foci were comparable between *Atf7ip2*^+/-^ and *Atf7ip2*^-/-^ H1T-positive mid/late pachytene spermatocytes (Fig. 3D). While a recent study of a separate *Atf7ip2*^-/-^ mouse line reported reduced numbers of XY pseudoautosomal regions (PARs) with MLH1 foci (Shao et al. 2023), our observations showed no significant difference in the proportions of MLH1-associated PARs in *Atf7ip2^+/-^* and *Atf7ip2^-/-^*models (Fig. 3D). Next, we analyzed chromosome synapsis by immunostaining for SYCP3 (a marker of both unsynapsed and synapsed axes) and SYCP1 (a marker of only synapsed axes); we observed occasional but significant autosomal asynapsis in *Atf7ip2*^-/-^ pachytene spermatocytes: ∼87% of *Atf7ip2*^-/-^ pachytene nuclei evidenced complete synapsis, while nearly all *Atf7ip2*^+/+^ spermatocytes showed complete synapsis (Fig. 3E). On occasion, the shapes of sex chromosome axes exhibited abnormal configurations, including apparent looped synapsis (“bubbles”), synapsis with large portions of itself (“irregular”), and synapsis at ends (“circular;” Fig. 3F); ∼25% of *Atf7ip2*^-/-^ pachytene nuclei demonstrated abnormal sex chromosome synapsis (Fig. 3F). These results suggest that, although ATF7IP2 may not play an outsized role in meiotic recombination, both DDR signaling and chromosome synapsis are impaired to some extent in *Atf7ip2*^-/-^ spermatocytes.

### ATF7IP2 directs SETDB1 and H3K9 methylation in male meiosis

Because ATF7IP binds SETDB1 to regulate H3K9me3 in somatic cells (Ichimura et al. 2005), we suspected that ATF7IP2 regulates H3K9me3 during meiosis. In meiotic prophase I, H3K9me3 accumulates on autosomal PCH and the sex chromosomes, where it is subject to dynamic regulation as XY undergoes MSCI (van der Heijden et al. 2007); H3K9me3 on the sex chromosomes is established by the methyltransferase SETDB1 (Hirota et al. 2018; Abe et al. 2022). Consistent with this, we observed normal H3K9me3 accumulation on autosomal PCH and XY chromatin in wild-type pachytene nuclei (Fig. 4A). Through careful examination, we noted multiple H3K9me3 accumulation patterns on the sex chromosome in the early pachytene stage of wild-type spermatocytes, coming to recognize four general patterns: Class I, covering the entirety of XY; Class II, covering the entirety of Y and X-PCH; Class III, covering X-PCH only; and Class IV, absent from XY, i.e., no signal (Fig. 4A, B; Supplemental Fig. S4A). We evaluated the proportions of patterns, finding that H3K9me3 enrichment on the XY domain was impaired in the early pachytene stage of *Atf7ip2*^-/-^ spermatocytes: 33% of nuclei showed essentially no signal anywhere on the XY chromosomes (Class IV), a pattern that was not observed in any *Atf7ip2*^+/+^ early pachytene nuclei (Fig. 4A, B). In normal mid and late pachytene stages, H3K9me3 is retained on X-PCH as it disappears from the remainder of XY chromatin (Fig. 4A), presumably due to histone replacement and the incorporation of histone variant H3.3 (van der Heijden et al. 2007). Then, in the normal diplotene stage, H3K9me3 signals promulgate through the XY chromatin in a likely reflection of H3K9me3’s *de novo* deposition on H3.3 (Fig. 4A). However, in *Atf7ip2*^-/-^ mid and late pachytene spermatocytes, H3K9me3 on X-PCH decreased, and reestablishment through the entirety of XY did not take place in the diplotene stage (Fig. 4A, C). Alongside the diplotene reestablishment of H3K9me3, H3K9me2 also accumulates on XY chromatin; however, in *Atf7ip2*^-/-^ diplotene spermatocytes, we noted a clear loss of H3K9me2 (Fig. 4D, E). Concomitant with these changes, in *Atf7ip2^-/-^*spermatocytes, we observed the strong accumulation of H3K9 acetylation (H3K9ac), which counteracts H3K9 methylation, on X-PCH (Supplemental Fig. S4B). On the other hand, proportions of H3K9me1 accumulation patterns were unchanged between control and *Atf7ip2*^-/-^ spermatocytes (Supplemental Fig. S4C, D), highlighting a specific role for ATF7IP2 in the regulation of H3K9me2/3 and H3K9ac. Together, these results indicate that ATF7IP2 is required for the establishment of H3K9me2/3 on diplotene XY chromatin, consistent with the concurrent, dynamic localization of ATF7IP2 from X-PCH through XY chromatin.

**Figure 4:**
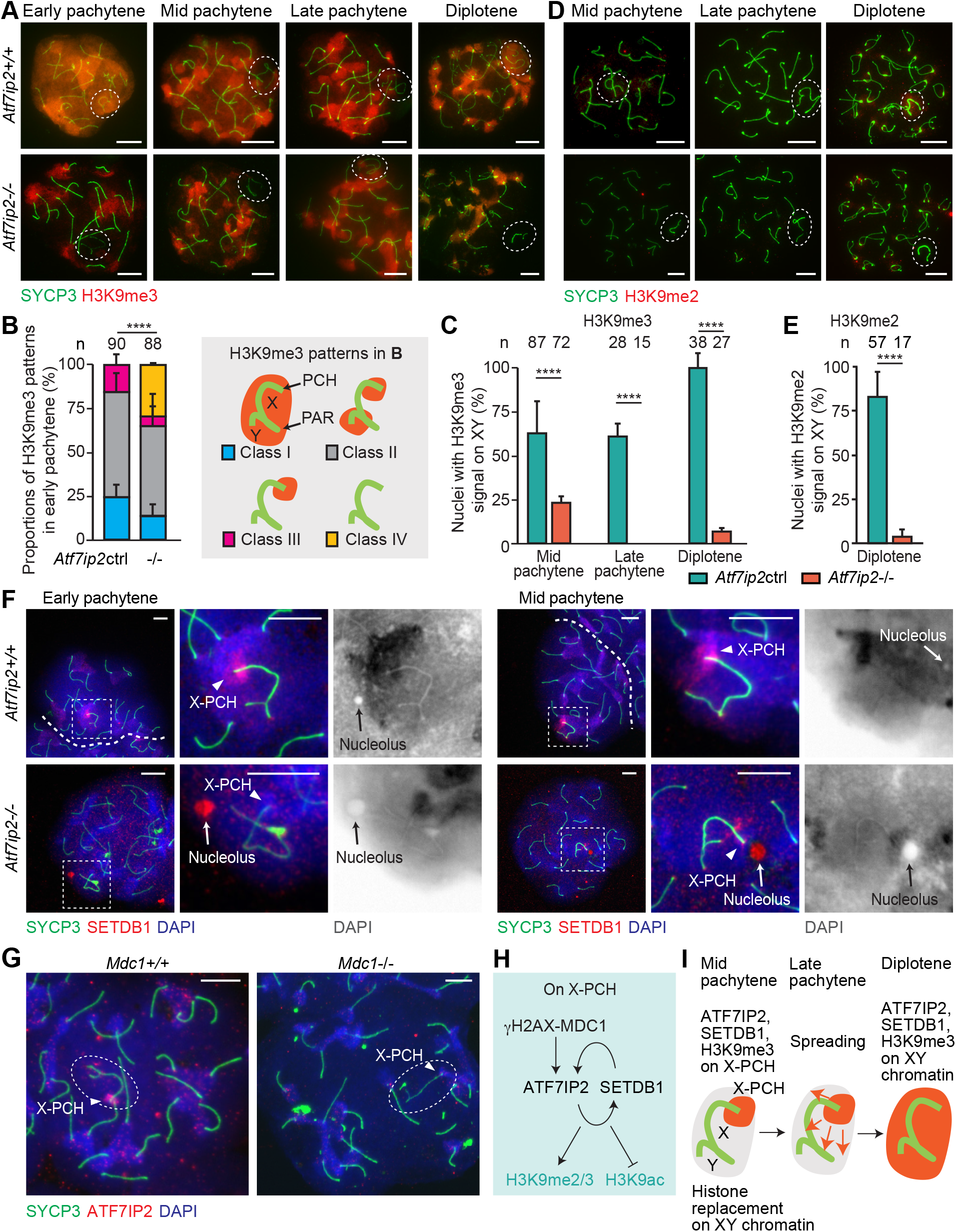
ATF7IP2 is required for H3K9 methylation on the sex chromosomes during male meiosis. **(A)** *Atf7ip2^+/+^* and *Atf7ip2^-/-^* spermatocyte chromosome spreads stained with antibodies raised against H3K9me3 and SYCP3 (a marker of chromosome axes, both synapsed and unsynapsed). Dashed circles indicate the sex chromosomes. Scale bars: 10 μm. **(B)** H3K9me3 accumulation patterns on the sex chromosomes of *Atf7ip2^+/+^* and *Atf7ip2^-/-^*early pachytene spermatocytes. Patterns are classified with four criteria (see right). Three independent experiments. P-values are from Fisher’s exact tests: **** < 0.0001. Scale bars: 10 μm. **(C)** Quantification of mid pachytene, late pachytene, and diplotene spermatocytes with H3K9me3 signals on the sex chromosomes. Three independent experiments. P-values are from Fisher’s exact tests: * < 0.05, *** < 0.001, **** < 0.0001. **(D)** Chromosome spreads stained with antibodies raised against H3K9me2 and SYCP3. **(E)** Quantification of diplotene spermatocytes with H3K9me2 signals on the sex chromosomes. Three independent experiments. P-values are from Fisher’s exact tests, **** < 0.0001. **(F)** Chromosome spreads stained with antibodies raised against SETDB1 and SYCP3. Dashed squares are magnified in the panels to the right. Scale bars: 10 μm. **(G)** *Mdc1^+/+^* and *Mdc1^-/-^* spermatocyte chromosome spreads stained with antibodies raised against ATF7IP2 and SYCP3. Scale bars: 10 μm. **(H)** Summary of the γH2AX/MDC1-ATF7IP2-SETDB1 pathway on X-PCH. **(I)** Schematic: establishment of H3K9me3 on the sex chromosomes in normal mid pachytene-to-diplotene spermatocytes.

Given this, we hypothesized that ATF7IP2 regulates the spatiotemporal recruitment of SETDB1, which mediates H3K9me3, to the sex chromosomes. In wild-type early and mid pachytene nuclei, SETDB1 localizes on the XY chromosomes and is notably enriched on the X-PCH. In corresponding *Atf7ip2^-/-^*spermatocytes, SETDB1 was not enriched on X-PCH, localizing instead to sex chromosome-adjacent nucleoli (Fig. 4F). Consistent with this, *Atf7ip2*^-/-^ X-PCH was less DAPI-intense compared to controls (Supplemental Fig. S5), suggesting a defect in heterochromatin formation. We also noticed that the pachytene accumulation of SETDB1 on autosomal PCH was disrupted in corresponding *Atf7ip2*^-/-^ spermatocytes: In contrast to the constrained, intense SETDB1 signals of wild-type samples, we observed diffuse SETDB1 signals through the whole of mutant nuclei (Fig. 4F). We also observed that, in *Setdb1* conditionally deleted mutants driven by the germline-specific *Ddx4*-Cre (*Setdb1*-cKO) (Abe et al. 2022), the accumulation of ATF7IP2 on X-PCH was significantly reduced (Supplemental Fig. S6A). Taken together, these results indicate a pan-nuclear role for ATF7IP2 in the spatiotemporal regulation of SETDB1; furthermore, at X-PCH, ATFIP2 and SETDB1 likely operate in tandem, possibly as a protein complex.

A hallmark of normal MSCI is the sex chromosome-wide accumulation of γH2AX, and γH2AX domain formation is tightly associated with the initiation and maintenance of MSCI (Fernandez-Capetillo et al. 2003; Abe et al. 2022). γH2AX domain formation is directed by MDC1, a γH2AX-binding protein and central mediator of the DDR, through a feed-forward mechanism (Ichijima et al. 2011). We hypothesized that the accumulation of ATF7IP2 on XY chromatin occurs downstream of MDC1. To test this, we stained for ATF7IP2 in *Mdc1*^-/-^ spermatocytes, finding that, in the absence of MDC1, ATF7IP2 failed to concentrate on X-PCH (Fig. 4G). Similarly, the accumulation of SETDB1 on X-PCH depended on MDC1 (Supplemental Fig. S6B). These results suggest that the MDC1-dependent DDR pathway regulates ATF7IP2 and SETDB1 localization on the sex chromosomes. We infer that, in pachytene spermatocytes, the MDC1-dependent DDR pathway recruits ATF7IP2, and thus SETDB1, to X-PCH; in the subsequent diplotene stage, both factors spread through the XY chromatin and, as this occurs, SETDB1 deposits pan-XY H3K9me2/3 (Fig. 4H, I). Corroborating this model, we found that MDC1 accumulation on XY chromatin occurred independently of ATF7IP2 or SETDB1 (Supplemental Fig. S6C, D).

To parse mechanisms related to ATF7IP2, we tested the localization of related factors. A previous study suggested a role for the SETDB1-interacting protein TRIM28 as a linker between the DDR pathway and SETDB1 (Hirota et al. 2018). However, we found that TRIM28 does not localize on the sex chromosomes in wild-type meiosis (Supplemental Fig. S7A), which raises the possibility that ATF7IP2 works independently of TRIM28 to link the DDR pathway and SETDB1. In line with this possibility, TRIM28 is dispensable for male meiotic progression (Tan et al. 2020). Downstream of the DDR pathway, the chromatin remodeler CHD4 is recruited to X-PCH (Broering et al. 2014); in *Atf7ip2*^-/-^ spermatocytes, CHD4 accumulation on X-PCH was unchanged from controls (Supplemental Fig. S7B). The germline-specific Polycomb protein SCML2 also accumulates on XY chromatin downstream of the DDR pathway; in *Atf7ip2*^-/-^ spermatocytes, SCML2 localization did not differ from controls (Supplemental Fig. S7C, D). These results suggest that the disfunction of ATF7IP2 is not related to the localization of TRIM28, CHD4, and SCML2.

We also examined the localization of ATF7IP in *Atf7ip2*^-/-^ spermatogenesis. In wild-type tissue sections, ATF7IP was predominantly found in the nuclei of primary spermatocytes (Supplemental Fig. S7E); more specifically, it localized to the X-PCH in wild-type pachytene spermatocytes (Supplemental Fig. S7F, G). Contrastingly, in *Atf7ip2*^-/-^ pachytene spermatocytes, ATF7IP was absent from the X-PCH and, instead, localized to the XY PAR. These results demonstrate ATF7IP2 is essential for directing ATF7IP and SETDB1 to X-PCH in pachytene spermatocytes.

### ATF7IP2 is required for meiotic gene regulation

Having established its meiotic phenotype and essential role in H3K9 methylation, we sought to investigate the function of ATF7IP2 in meiotic gene regulation. To this end, we performed single-cell RNA sequencing (scRNA-seq) analyses of whole testicular cells from *Atf7ip2*^-/-^ mice and their *Atf7ip2*^+/+^ littermates at P15. The cellular composition of the testis changes as development progresses, leading us to confirm that, in P15 testes, the first wave of spermatogenesis exhibited a similar cellular composition between *Atf7ip2*^+/+^ and *Atf7ip2*^-/-^ mice. Indeed, we observed this was the case until the mid-to-late pachytene stages, when defects appeared based on immunostaining against major markers of spermatogenesis, including ZBTB16, STRA8, SYCP3, γH2AX, and H1T (Supplemental Fig. S8).

Using the scRNA-seq data, we endeavored to determine when ATF7IP2 functions in wild-type spermatogenesis. Since ATF7IP2 expression was restricted to germ cells, scRNA-seq data derived from germ cell populations (spermatogonia and spermatocytes) were analyzed apart from those of testicular somatic cells (Sertoli cells, Leydig cells, peritubular myoid cells, endothelial cells, and hemocytes) (Fig. 5A; Supplemental Fig. S9A, B). Using the UMAP of scRNA-seq data from *Atf7ip2*^+/+^ and *Atf7ip2*^-/-^ germ cell populations, we identified 13 cell-type clusters; the cluster numbers are based on the numbers of cells comprising each cluster: Cluster 0 is the largest, and Cluster 12 is the smallest (Fig. 5B, C; Supplemental Fig. S9C, D). Assessing the expression of key marker genes for spermatogenesis with respect to the UMAP, we inferred the developmental trajectory of P15 spermatogenesis (Fig. 5D). As suggested by the high expression of *Gfra1*, Cluster 8 represented a population of undifferentiated spermatogonia, including spermatogonial stem cells. Cluster 1 represented a population of differentiating spermatogonia as indicated by the initial upregulation of *Stra8.* Clusters 6 and 7 represented cells at the initiation of meiosis, consistent with the upregulation of *Meiosin* and *Stra8.* Cluster 11 represented a population of spermatocytes in early meiotic prophase as denoted by the upregulation of *Prdm9.* Based on the expression of marker genes with respect to the UMAP, we inferred that spermatogenesis progressed along the trajectory from Clusters 8 to 12. Although *Atf7ip2* was expressed in a broad range of spermatogenic stages, its expression level was higher in Clusters 7, 6, 11, and 5, all of which correspond to meiotic prophase (Fig. 5E). Given that *Atf7ip2* is bound by MEIOSIN and STRA8 (Fig. 1B), it is possible that the expression of *Atf7ip2* was boosted, rather than initiated, at the entry to meiosis. In contrast, *Atf7ip* is constitutively expressed in spermatogenesis (Fig. 5E).

**Figure 5.**
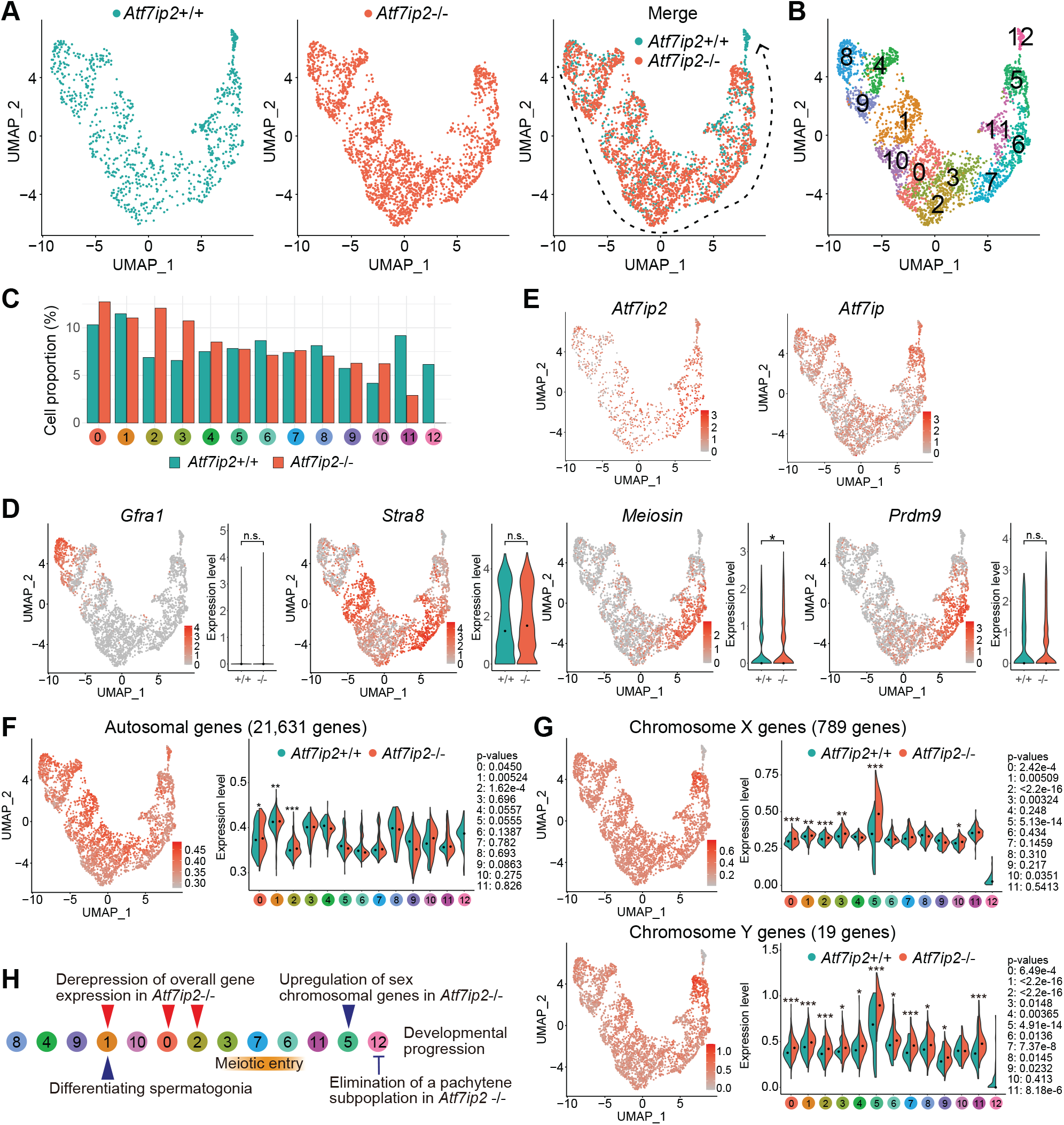
scRNA-seq analyses of *Atf7ip2*^+/+^ and *Atf7ip2*^-/-^ spermatogenic germ cells. **(A)** UMAP representations of scRNA-seq transcriptome profiles for germ cells from *Atf7ip2*^+/+^ testes (left: P15), *Atf7ip2*^-/-^ testes (middle: P15), and both *Atf7ip2*^+/+^ and *Atf7ip2*^-/-^ testes (right). Gray arrow: inferred developmental trajectory. **(B)** Clustering of UMAP-projected scRNA-seq transcriptome profiles for *Atf7ip2*^+/+^ and *Atf7ip2*^-/-^ germ cells based on gene expression patterns. **(C)** Bar graph showing the proportions of *Atf7ip2*^+/+^ and *Atf7ip2*^-/-^ germ cells among the clusters. **(D)** UMAP representations showing expression patterns for key developmental marker genes in spermatogenic cells. Genes include *Gfra1*, which represent undifferentiated spermatogonia; *Stra8*, differentiating spermatogonia; *Meiosin*, preleptotene spermatocytes; and *Prdm9*, early meiotic prophase spermatocytes. P-values are from Wilcoxon rank sum tests: n.s., not significant; * < 0.05. **(E)** Expression patterns for *Atf7ip2* and *Atf7ip* upon the UMAP. **(F)** Expression levels for autosomal genes. P-values are from Wilcoxon rank sum tests: * < 0.05, ** < 0.01, *** < 0.001. **(G)** Expression levels for X chromosomal genes (top) and Y chromosomal genes (bottom). P-values are from Wilcoxon rank sum tests: * < 0.05, ** < 0.01, *** < 0.001. **(H)** Summary of *Atf7ip2*^-/-^ phenotypes in spermatogenic germ cells. Subtype clusters are ordered by inferred developmental progression. Key cell types and events in *Atf7ip2^+/+^* and *Atf7ip2^-/-^*spermatogenesis are shown.

Next, we sought to understand when cell death takes place in *Atf7ip2*^-/-^ spermatocytes. Starting from Cluster 8 through to Cluster 5, the gene expression profiles for *Atf7ip2*^+/+^ and *Atf7ip2*^-/-^ germ cell populations overlapped one another to a high degree (Fig. 5A, B), indicating that *Atf7ip2*^-/-^ spermatogenesis progressed until the stage of spermatogenesis that corresponds to Cluster 5. However, we noticed certain subpopulations—Clusters 10, 0, 2, and 3, representing B spermatogonia through to preleptotene cells—were more numerous in *Atf7ip2*^-/-^ germ cells (Fig. 5C); the increased cluster sizes suggest that, in the absence of ATF7IP2, the entry into meiosis is hampered. Furthermore, the subpopulation represented by Cluster 12 was present in *Atf7ip2*^+/+^ germ cells but missing amid *Atf7ip2*^-/-^ germ cells (Fig. 5B, C). Furthermore, in *Atf7ip2*^+/+^ germ cells, expression levels of sex-linked genes were abruptly downregulated in the transition from Clusters 5 to 12 (Fig. 5G). Intriguingly, in *Atf7ip2*^-/-^ cells, Cluster 5 was associated with a strong, abrupt upregulation of sex-linked gene expression (Fig. 5G), suggesting that MSCI failure began in the Cluster-5 subpopulation of *Atf7ip2*^-/-^ cells. Thus, the loss of the Cluster-12 subpopulation in *Atf7ip2*^-/-^ testes was preceded by an ectopic upregulation of X and Y chromosomal genes in Cluster 5 (Fig. 5G), indicating Clusters 5 and 12 represent pachytene spermatocytes.

Remarkably, gene enrichment analysis revealed that genes related to late spermatogenesis (e.g., *Clgn, Hspa2, Piwil1,* and *Ldhc*) were highly expressed in the Cluster-12 subpopulation of spermatocytes (Supplemental Fig. S9C, Table S1). Since those genes are known to be expressed in the late pachytene stage onward, we infer Cluster 12 corresponds to cytologically defined late pachytene spermatocytes. This is consistent with the cytological observation that *Atf7ip2*^-/-^ spermatocytes progressed through early meiotic prophase but were eliminated via apoptosis at the transition from the late pachytene to diplotene stages (Fig. 3B). Thus, ATF7IP2 is required for spermatocytes to progress beyond the late pachytene stage represented by Cluster 12.

### ATF7IP2 binds broadly to the sex chromosomes and autosomal gene promoters

To determine where ATF7IP2 binds the genome of wild-type pachytene spermatocytes, we performed CUT&Tag for ATF7IP2 in two biological replicates. The replicates were highly correlated (Supplemental Fig. S10A), allowing us to merge them for downstream analyses. Analyses of ATF7IP2 coverage revealed 61,797 genome-wide regions of enriched ATF7IP2-binding, i.e., “ATF7IP2 peaks” (Fig. 6A). We observed ATF7IP2 peaks on TSSs (26.7 %), gene bodies (27.7 %), and intergenic regions (45.4 %). TSS peaks were enriched on autosomes, while intergenic peaks were enriched on the sex chromosomes (Fig. 6A), suggesting distinct functions for ATF7IP2 on the autosomes and sex chromosomes. Continuing to analyze wild-type pachytene spermatocytes, we performed two-step clustering with ATF7IP2 peaks, regions of H3K9me3 coverage, and regions of coverage for the active promoter mark H3K4me3, generating three clusters (Fig. 6B). Cluster I regions (6,632) are associated with H3K4me3 deposition (Fig. 6B) on autosomes and at TSSs (Fig. 6C, D). Cluster II regions (22,579) are associated with broad H3K9me3 enrichment (Fig. 6B); 70% of these regions are on the sex chromosomes (Fig. 6C), mostly at intergenic regions and gene bodies (Fig. 6D); this is in line with the role of ATF7IP2 in the regulation of H3K9me3 on the sex chromosomes. Cluster III regions (32,628) largely represent autosomal intergenic regions and gene bodies (Fig. 6C, D).

**Figure 6.**
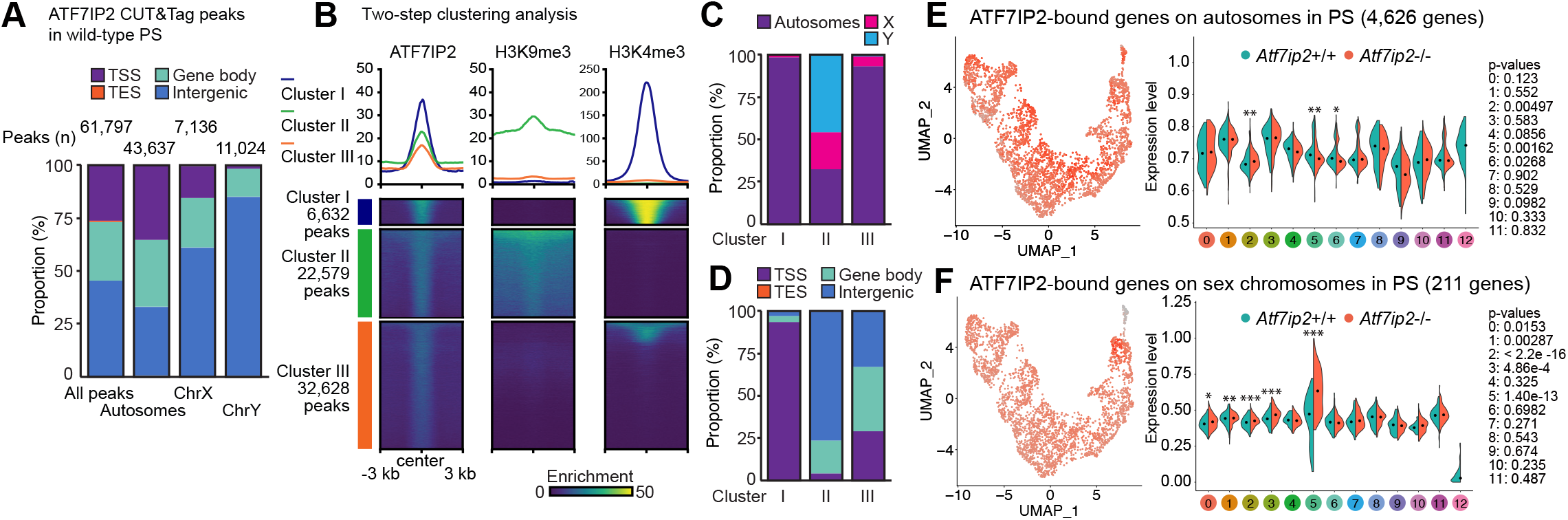
ATF7IP2-binding sites in pachytene spermatocytes. **(A)** Numbers and genomic distribution of ATF7IP2 CUT&Tag peaks in wild-type pachytene spermatocytes. **(B)** Two-step clustering analysis of ATF7IP2 CUT&Tag peaks and H3K9me3 and H3K4me3 enriched-regions. Average tag density profiles (top) and heatmaps for each cluster (bottom). **(C)** Chromosomal distribution of ATF7IP2 peak clusters. **(D)** Genomic distribution of ATF7IP2 peak clusters. **(E)** Expression levels of ATF7IP2-bound autosomal genes in scRNA-seq. P-values are from Wilcoxon rank sum tests: * < 0.05, ** < 0.01. **(F)** Expression levels for ATF7IP2-bound sex chromosomal genes in scRNA-seq. P-values are from Wilcoxon rank sum tests: * < 0.05, ** < 0.01, *** < 0.001

Based on the enrichment of ATF7IP2 at TSSs, we sought to identify ATF7IP2-target genes in pachytene spermatocytes. Our analyses revealed 4,917 autosomal genes and 270 sex chromosomal genes (Supplemental Table S2). ATF7IP2 binds the promoters of a broad range of genes required for meiotic prophase and spermiogenesis, including *Hormad1* and *Sycp3*, both autosomal genes, as well as Y-linked *Zfy1.* These promoter peaks are associated with the active histone modifications H3K4me3 and H3K27ac (Supplemental Fig. S10B). Next, using our scRNA-seq data set for *Atf7ip2*^+/+^ and *Atf7ip2*^-/-^ pachytene spermatocytes, we sought to understand the regulation of ATF7IP2-target genes. We detected the expression of 4,626 ATF7IP2-target genes on autosomes and 211 on the sex chromosomes. The autosomal genes were downregulated in *Atf7ip2*^-/-^ pachytene spermatocytes (Fig. 6E: Clusters 6 to 5, representing the early-to-mid-pachytene stages). Because wild-type autosomal promoter peaks are associated with H3K4me3, these results indicate that ATF7IP2 binds to and positively regulates the expression of these genes. On the other hand, in *Atf7ip2*^-/-^ pachytene spermatocytes, the 211 sex chromosomal genes are highly upregulated in Cluster 5 (mid pachytene spermatocytes), indicating that ATF7IP2 binds to and negatively regulates the expression of these genes. These results reveal two separate functions for ATF7IP2 in pachytene spermatocytes: one for autosomal gene expression and, contrastingly, another for sex chromosomal gene repression.

### ATF7IP2 directs meiotic gene regulation

To elucidate gene regulatory mechanisms associated with ATF7IP2, we isolated pachytene spermatocytes from *Atf7ip2*^+/+^ and *Atf7ip2*^-/-^ testes, verified their purity (Supplemental Fig. S10D), performed bulk RNA-seq with spike-in controls, and analyzed the resulting transcription data. In isolating the cells, we were surprised to observe that *Atf7ip2*^-/-^ spermatocytes were smaller than their *Atf7ip2*^+/+^ counterparts (Supplemental Fig. S10D). This gross decrease in size suggests that, in *Atf7ip2*^-/-^ spermatocytes, the pachytene transcriptional burst (Maezawa et al. 2020) is compromised—a possibility consistent with the global downregulation of ATF7IP2-bound autosomal genes detected with scRNA-seq.

Comparing the spike-in-normalized mutant and control RNA-seq data, we identified 8,507 autosomal differentially expressed genes (DEGs): 185 upregulated and 8,322 downregulated (Fig. 7A). To understand how autosomal DEGs are expressed during normal spermatogenesis, we reanalyzed separate RNA-seq data taken from cell types sampled across wild-type spermatogenesis (Maezawa et al. 2018b). The 185 upregulated genes displayed high expression levels in wild-type spermatogonia but were suppressed in wild-type pachytene spermatocytes (Supplemental Fig. S11A). Thus, their upregulation in *Atf7ip2*^-/-^ pachytene spermatocytes suggests an ectopic expression of normally repressed premeiotic genes. The top Gene Ontology (GO) (Ashburner et al. 2000) enrichment terms for these genes are related to immune-system functions (Supplemental Fig. S11B), suggesting that ATF7IP2 suppresses the expression of immune genes in pachytene spermatocytes. In contrast, many of the 8,322 downregulated genes were highly expressed in wild-type pachytene spermatocytes (Supplemental Fig. Fig. S11A), and the associated GO enrichment terms were related to spermatogenesis (Supplemental Fig. S11B). These findings indicate that many spermatogenesis-related genes fail to activate in *Atf7ip2*^-/-^ pachytene spermatocytes. Shifting focus to the sex chromosomes, we detected 528 DEGs associated with the *Atf7ip2*^-/-^ pachytene X chromosome: 522 were upregulated in mutants relative to controls, while 6 were downregulated (Fig. 7A). On the Y chromosome, 12 DEGs were upregulated, and we detected no downregulated DEGs (Fig. 7A). These results are largely consistent with bulk RNA-seq analyses of P14 juvenile testes (Supplemental Fig. S12), together indicating that MSCI is disrupted in *Atf7ip2^-/-^*pachytene spermatocytes.

**Figure 7.**
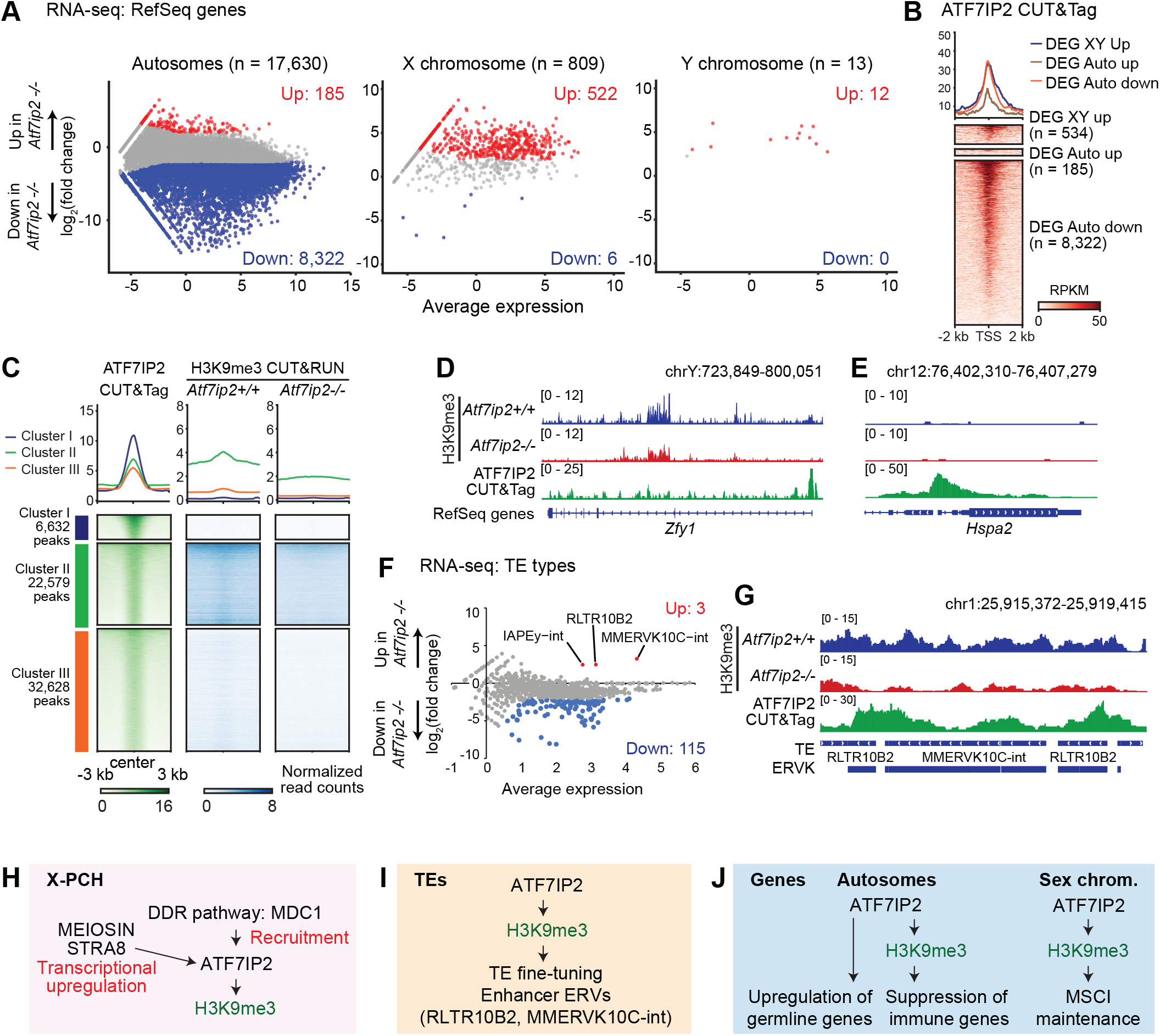
ATF7IP2 directs meiotic gene regulation and regulates TEs. **(A)** Comparison of *Atf7ip2*^+/+^ and *Atf7ip2*^-/-^ pachytene spermatocyte transcriptomes. Autosomal, X, and Y genes were analyzed separately. Two independent biological replicates were examined. All genes with adjusted p-values (Benjamini-Hochberg method) are plotted. Differentially expressed genes (DEGs: log_2_ fold change ≥ 2, adjusted p-value ≤ 0.05) are colored (red: upregulated in *Atf7ip2*^-/-^ testes; blue: downregulated in *Atf7ip2*^-/-^ testes), and numbers are shown. **(B)** ATF7IP2 CUT&Tag enrichment at DEG TSSs ± 2 kb in pachytene spermatocytes isolated from *Atf7ip2*^-/-^ mice. Average tag density profiles (top) and heatmaps for each cluster (bottom). **(C)** ATF7IP2 CUT&Tag and H3K9me3 CUT&RUN enrichment in Clusters I–III (defined in Fig. 6B). Average tag density profiles (top) and heatmaps for each cluster (bottom). **(D, E)** Track views of the *Zfy1* locus (an upregulated Y-linked locus) and the *Hspa2* locus (a downregulated autosomal locus). **(F)** Comparison of *Atf7ip2*^+/+^ and *Atf7ip2*^-/-^ pachytene spermatocyte transposable element (TE) expression. All TE types are plotted. Differentially expressed TE types (DEGs: log_2_ fold change > 2, adjusted p-value < 0.05) are colored (red: upregulated in *Atf7ip2*^-/-^; blue: downregulated in *Atf7ip2*^-/-^), and numbers are shown. **(G)** Track view of the ATF7IP2-targeted TEs RLTR10B2 and MMERVK10C-int. **(H)** Summary and model of the function of ATF7IP2 on X-PCH. **(I)** Summary and model of the function of ATF7IP2 in TE regulation. **(J)** Summary and model of the function of ATF7IP2 in gene expression regulation.

MSCI is initiated by the DDR pathway and maintained through active DDR signaling (Ichijima et al. 2011; Abe et al. 2022). On the *Atf7ip2^-/-^*XY domain, γH2AX signals were observed (Fig. 2), but H3K9me2/3 deposition was not established as the pachytene stage progressed into the diplotene stage (Fig. 4). Indeed, in place of H3K9me2/3, we observed signals for the active transcriptional mark H3K9ac (Supplemental Figure S4B). Thus, we suspect that MSCI is initiated but not maintained in *Atf7ip2*^-/-^ spermatocytes. To test this, we stained for POLII in *Atf7ip2*^-/-^ pachytene spermatocytes. In normal mid pachytene spermatocytes, we observed the exclusion of POLII from XY domains in 100% of observed nuclei (n = 65, Supplemental Fig. S11C), confirming the accurate detection of MSCI through POLII immunostaining. However, in *Atf7ip2*^-/-^ mid pachytene spermatocytes, we observed the exclusion of POLII from XY chromatin in only 73.3% of nuclei (n = 105, Supplemental Fig. S11D); 26.7% of *Atf7ip2*^-/-^ nuclei saw the inclusion of POLII in XY domains (Supplemental Fig. S11E)— evidence for defective MSCI. These results suggest that, in the absence of ATF7IP2, the initiation of MSCI occurs, but MSCI fails to be maintained.

In terms of γH2AX signals on XY chromatin and the loss of H3K9me3 deposition, the *Atf7ip2*^-/-^ phenotype overlaps the reported phenotype for *Setdb1-*cKO mice (Hirota et al. 2018; Cheng et al. 2021; Abe et al. 2022). To determine the relationship between *Atf7ip2* and *Setdb1* mutations, we reanalyzed *Setdb1-*cKO RNA-seq data for pachytene spermatocytes (Hirota et al. 2018) (Supplemental Fig. S12). Although MSCI was disrupted, the massive downregulation of autosomal genes was not observed in the *Setdb1-*cKO spermatocytes. Thus, we infer that ATF7IP2’s gene regulatory functions are broader in consequence than those of SETDB1.

To understand the mechanism through which ATF7IP2 regulates H3K9me3 deposition, we produced and analyzed H3K9me3 CUT&RUN data from *Atf7ip2*^+/+^ and *Atf7ip2*^-/-^ pachytene spermatocytes (Supplemental Fig. 10F). We found that H3K9me3 is largely dependent on ATF7IP2, especially at the sites of Cluster II ATF7IP2-bound peaks (Fig. 7C; Cluster II peaks were defined in Fig. 6B). ATF7IP2 and H3K9me3 signals frequently overlapped, with many regions of H3K9me3 deposition centered on ATF7IP2 peaks (Fig. 7C). Notably, H3K9me3 deposition was completely absent or strongly diminished in the *Atf7ip2^-/-^* model, indicating H3K9me3 enrichment is dependent on ATF7IP2 (Fig. 7C). As shown in a track view of the Y-linked *Zfy1* locus, ATF7IP2-binding sites frequently align with, or are immediately adjacent to, locations of ATF7IP2-dependent H3K9me3 (Fig. 7D). Conversely, there was no observed H3K9me3 enrichment on ATF7IP2-dependent autosomal genes, as evidenced by loci such as *Hspa2* (Fig. 7E). We conclude that ATF7IP2 directs H3K9me3 deposition while simultaneously orchestrating meiotic gene activation on autosomes, much of which is independent of H3K9me3.

### ATF7IP2 fine-tunes the expression of transposable elements

SETDB1-mediated H3K9me3 is a well-known suppressor of transposable elements (TEs) (Matsui et al. 2010; Rowe et al. 2013). Therefore, we sought to examine TE expression using our RNA-seq data in combination with a “best-match” TE annotation set (Sakashita et al. 2020), which enables the detection of alignments uniquely mapped to TEs that are not exon-derived (mRNA-derived). This strategy eliminates detection of TEs that are parts of mRNA, preventing the conflation of mRNA and TE expression. In *Atf7ip2*^-/-^ pachytene spermatocytes, three TE types (IAPEy-int, RLTR10B2, MMERVK10C-int) were upregulated, while 115 types were downregulated (Fig. 7F). ATF7IP2 bound these upregulated TEs, and H3K9me3 at these loci was ATF7IP2-dependent (Fig. 7G). In wild-type spermatogenesis, TE expression undergoes dynamic changes at the mitosis-to-meiosis transition, and a subset of TEs—specifically endogenous retrovirus K (ERV) families that are also known as long-terminal repeats (LTRs)— are activated as enhancers in meiosis (Sakashita et al. 2020). Notably, these meiotic enhancer ERVs (RLTR10B2) are among the upregulated TEs (Fig. 7F). The meiotic enhancer ERVs are active in wild-type pachytene spermatocytes and were further upregulated in *Atf7ip2*^-/-^ cells. Thus, ATF7IP2 may fine-tune the activity of these TEs. In contrast, various TE types, particularly those enriched with LTRs and active in the pachytene stage, were downregulated in *Atf7ip2*^-/-^ spermatocytes. In all, our study identifies distinct functions for ATF7IP2 in regulating protein-coding genes on autosomes and sex chromosomes, as well as in the regulation of TEs (Fig. 7H, I, J).

## Discussion

Our study identifies ATF7IP2 as a counterpart to ATF7IP that is highly expressed in the male germline and directs SETDB1-mediated H3K9 methylation—a conclusion supported by two major observations. First, in wild-type meiosis, ATF7IP2, SETDB1, and H3K9me3 accumulate on autosomal PCH; in pachytene spermatocytes, all are enriched on the X-PCH, the site from which they spread through the diplotene XY domain. Second, in *Atf7ip2*^-/-^ pachytene spermatocytes, SETDB1 was grossly delocalized, and H3K9me2/3 was not present on the XY chromatin in the late pachytene-to-diplotene stages. As might be expected, the *Atf7ip2*^-/-^ meiotic phenotype overlaps to some extent the meiotic phenotype of *Setdb1-*cKO mice. Thus, our study reveals the molecular logic for the management of SETDB1 and H3K9me3 in meiosis, demonstrating the unique nature of meiotic heterochromatin and its distinct regulation with respect to autosomes and the sex chromosomes.

However, in the early pachytene stage, there is a phenotypic difference between *Atf7ip2*^-/-^ and *Setdb1-*cKO mice with regards to H3K9me3 localization: H3K9me3, a SETDB1-dependent marker of XY chromatin (Hirota et al. 2018; Abe et al. 2022), is affected but not completely absent from XY chromatin in *Atf7ip2*^-/-^ mice. Thus, there may be an alternate regulator of SETDB1 in early pachytene spermatocytes. In support of this possibility, *Setdb1-*cKO spermatocytes evidenced more severe chromosome synapsis defects (Hirota et al. 2018; Cheng et al. 2021; Abe et al. 2022) than *Atf7ip2*^-/-^ spermatocytes. Nevertheless, the meiotic arrest phenotype indicates that the ATF7IP2-dependent regulation of SETDB1 (likely through an ATF7IP2-SETDB1 complex) and H3K9me3 becomes essential in the pachytene-to-diplotene transition.

Our study also reveals novel aspects of the meiotic sex chromosomes. We propose that, through the recruitment of SETDB1, ATF7IP2 functions as an effector that links DDR signaling and SETDB1-mediated H3K9me3. The γH2AX-binding partner MDC1 is necessary for the recruitment of ATF7IP2 to X-PCH (Fig. 4G). A previous study proposed that TRIM28 (KAP1)—a SETDB1 partner in ERV suppression—links the DDR and SETDB1 on the meiotic sex chromosomes (Hirota et al. 2018). However, we did not observe TRIM28 enrichment on XY, and so we question TRIM28’s status as a linker. Furthermore, it was reported that young *Trim28* mutant mice are initially fertile and only become sterile with age (Tan et al. 2020), indicating that TRIM28 is not essential for MSCI.

We find that the establishment of H3K9me2/3 on diplotene XY chromatin is ATF7IP2-dependent. Given the extensive histone replacement that occurs in MSCI (H3.1/H3.2 to H3.3) (van der Heijden et al. 2007), H3K9me2/3 deposition is likely to take place on “fresh” H3.3 in a process that is also ATF7IP2-dependent. In the latter stages of spermatogenesis, H3K9me2/3 is a persistent mark on the sex chromosomes, from MSCI to postmitotic silencing (Namekawa et al. 2007); thus, the ATF7IP2-dependent mechanisms described here could be driving heritable epigenetic states through meiotic divisions.

Unexpectedly, our study demonstrates that ATF7IP2 is required for global gene regulation in pachytene spermatocytes. In mid pachytene spermatocytes, a burst of gene activation takes place, and this is driven by the transcription factor A-MYB (MYBL1) through the activation of meiotic enhancers (Bolcun-Filas et al. 2011; Maezawa et al. 2020). Thus, there is an intriguing possibility that such meiosis-specific transcription requires ATF7IP2. Importantly, ATF7IP2 is present at thousands of autosomal promoters, where H3K9me3 is notably absent. Thus, ATF7IP2 could regulate transcriptional mechanisms independent of H3K9me3. Intriguingly, ATF7, an ATF7IP2-interacting protein, also accumulates on a wide range of autosomal promoters in testicular germ cells, mediating epigenetic inheritance through the regulation of H3K9me2 (Yoshida et al. 2020). In future studies, a key goal will be to determine the mechanistic relationship between ATF7IP2 and ATF7 in the context of meiotic gene regulation. While ATF7IP2’s localization on the sex chromosomes requires MDC1, it is unknown what regulates its recruitment to autosomes—although one possibility is ATM-dependent DDR signaling. Furthermore, it is unknown what coordinates ATF7IP2’s distinct autosomal and XY functions.

Finally, we show that ATF7IP2 fine-tunes the expression of retrotransposon-derived loci in male germ cells, a function that coincides with SETDB1’s role in TE silencing. In *Atf7ip2*^-/-^ spermatocytes, we observed the upregulated expression of immune genes, a phenomenon akin to SETDB1-mediated immune escape in tumorigenesis (Griffin et al. 2021). In tumor cells, the depletion of SETDB1 facilitates the expression of immune genes, thereby driving the intrinsic immunogenicity of tumors. Also in tumor cells, SETDB1 works together with the HUSH complex—itself functionally linked to ATF7IP (Timms et al. 2016)—to suppress large domains of the genome enriched for rapidly evolved TEs (Griffin et al. 2021). Notably, a large number of the germline genes activated in pachytene spermatocytes are rapidly evolved (Soumillon et al. 2013), as are the meiotic ERV enhancer loci that drive germline gene expression (Sakashita et al. 2020). In wild-type spermatocytes, these loci are associated with broad domains of H3K9me3. Furthermore, like many tumor cells, testicular germ cells are immunoprivileged, found beyond the blood-testes barrier. Given the similarities between germ and tumor cells, it is possible that ATF7IP2-directed SETDB1 mechanisms, which regulate MSCI and TEs, drive the quick-paced evolution of the germline genome. It may be that this work establishes a foundation to understand the mechanisms behind germline evolution in mammals.

A recent study reported another *Atf7ip2* mutant mouse line (Shao et al. 2023), and although the mouse phenotypes detailed in the two studies were largely consistent, we did not observe the reported difference in XY obligatory crossover (Shao et al. 2023). This could be due to an *Atf7ip2* mutational difference in the mouse lines. Further investigations are warranted to clarify the role of ATF7IP2 in male meiosis.

## Materials and Methods

### Animals

All mice were handled according to the guidelines of the Institutional Animal Care and Use Committee (IACUC: protocol no. IACUC2018-0040 and 21931) at Cincinnati Children’s Hospital Medical Center and the University of California, Davis.

### Generation of *Atf7ip2*^-/-^ mice

*Atf7ip2*^-/-^ mice were generated using a sgRNA (target sequence: TTCATGTCTACTCTTGCACT) that was selected according to location and the on- and off-target scores from the web tool CRISPOR (Haeussler et al. 2016).

### Preparation of meiotic chromosome spreads

Meiotic chromosome spread preparation, immunostaining, and data analysis were performed as described (Alavattam et al. 2018). Histology and immunostaining were performed as described (Abe et al. 2022).

### Isolation of pachytene spermatocytes

Isolation of pachytene spermatocytes using Fluorescence-activated cell sorting (FACS) was performed using SH800S (SONY), with Vybrant DyeCycle Violet Stain (DCV) (Invitrogen, V35003) stained testicular single-cell suspensions prepared as described previously (Yeh et al. 2021).

### Next-generation sequencing analysis

Library generation and data analyses for bulk RNA-seq, CUT&Tag, CUT&RUN, and scRNA-seq are described in the Supplemental Material. Other detailed experimental procedures are described in the Supplemental Material.

### Data Availability

RNA-seq data and CUT&RUN/Tag datasets were deposited in the Gene Expression Omnibus (accession: GSE244088). Testes bulk RNA-seq data reported in this paper were deposited in the Gene Expression Omnibus (accession: GSE223742). Single-cell RNA-seq data are available at DDBJ Sequence Read Archive (DRA) under the BioProject accession: PRJDB16643.

## Author contributions

K.G.A., J.M.E., M.H., R.S., K.-I.I., and S.H.N. designed the study. K.G.A., J.M.E., M.H., A.R.K., H.A., Y.K., Y.-H.Y., and J.K. performed experiments. K.G.A., J.M.E., M.H, A.R.K., H.A., M.H, Y.K. analyzed the mouse phenotypes. J.M.E., M.H isolated germ cells and performed scRNA-seq experiments. M.H performed bulk RNA-seq, CUT&Tag, CUT&RUN experiments. R.S. analyzed the scRNA-seq data. K.G.A., J.M.E., M.H, R.S., Y.M., K.O., S.Y., K.-I.I., and S.H.N. designed and interpreted the computational analyses. Y.-C.H. generated the *Atf7ip2*-/- mouse line. K.G.A., J.M.E., M.H., R.S., P.R.A, K.-I.I., and S.H.N. interpreted the results and wrote the manuscript with critical feedback from all other authors. S.H.N. supervised the project.

## Acknowledgments

We thank Yoshinori Watanabe, Hiroki Shibuya, Akihiro Morimoto, and Shohei Yamamoto for their contributions to the early stage of this investigation; members of the Namekawa laboratory for discussion and helpful comments regarding this manuscript; So Maezawa and Masashi Yukawa for the initial RNA-seq analysis; Neil Hunter and Richard M. Schultz for discussion; Yasuhiro Fujiwara and Yuki Okada for aiding in the transfer of the anti-ATF7IP2 antibody; the Transgenic Animal and Genome Editing Core at CCHMC for generation of the *Atf7ip2*^-/-^ mouse model; Yoichi Shinkai for providing the *Setdb1* floxed mouse line, Junjie Chen for providing the *Mdc1*-KO mice, and Mary Ann Handel for providing the anti-H1T antibody. Funding sources: NIH Training Program in Molecular and Cellular Biology T32GM007377 and Ford Foundation Predoctoral Fellowship to J.E.; R01 GM134731 to P.R.A.; KAKENHI #19H05743, #23H00379 and AMED PRIME #23gm6310021h0003, the program of the Research for High Depth Omics, IMEG, Kumamoto University to K.I.I.; UC Davis startup fund, and NIH R01 GM098605, R35 GM141085, and GM141085 diversity supplement to S.H.N.

