## Supplemental Material for "ATF7IP2/MCAF2 directs H3K9 methylation and meiotic gene regulation in the male germline"

### Methods

#### Animals

Mice were maintained on a 12:12 light:dark cycle in a temperature- and humidity-controlled vivarium (22 ± 2 °C; 40–50% humidity) with free access to food and water in a pathogen-free animal care facility. All mice were handled according to the guidelines of the Institutional Animal Care and Use Committee (IACUC: protocol no. IACUC2018-0040 and 21931) at Cincinnati Children's Hospital Medical Center and the University of California, Davis. At least three independent littermate pairs of different mice were used as biological replicates for all experiments, except for the next-generation sequencing experiments, which were performed using two independent biological replicates. C57BL/6 (hereafter described as B6) mice were used for all experiments using *Atf7ip2*$^{-/-}$ mice. To generate *Setdb1*-cKO (*Setdb1*$^{flox/flox}$; *Ddx4-Cre*$^{Tg/+}$) mice, a male carrying the *Setdb1*$^{flox/+}$ ; *Ddx4-Cre*$^{Tg/+}$ alleles was crossed with a female carrying the *Setdb1*$^{flox/flox}$ allele as described in our previous report (Abe et al. 2022). Males carrying the *Setdb1*$^{flox/+}$; *Ddx4-Cre*$^{Tg/+}$ alleles obtained from the same litter were used as littermate controls. Control and *Setdb1*cKO mice greater than 3 weeks of age were harvested and analyzed. The *Mdc1* knockout (*Mdc1*-KO) mice used in this study are described in our previous report (Ichijima et al. 2011). We define postnatal days (P) as days postpartum.

#### Generation of *Atf7ip2*$^{-/-}$ mice

*Atf7ip2*$^{-/-}$ mice were generated using a sgRNA (target sequence: TTCATGTCTACTCTTGCACT) that was selected according to location and on- and off-target scores from the web tool CRISPOR (Haeussler et al. 2016). The sgRNA was *in vitro* synthesized using a MEGAshorscript T7 kit (Thermo Fisher) and purified using a MEGAclear Kit (Thermo Fisher) according to the manufacturer's instructions. The sgRNA (100 ng/µl) and Cas9 protein (200 ng/µl, Thermo Fischer) were mixed at 37 °C for 15 min to form the ribonucleoprotein complex, then injected into the cytoplasm of C57BL/6N genetic-background one-cell-stage embryos by piezo-driven microinjection (Scott and Hu 2019). Injected embryos were immediately transferred into the oviductal ampulla of pseudopregnant CD-1 females. Live-born pups were genotyped by PCR and Sanger sequencing.

#### Preparation of meiotic chromosome spreads

Meiotic chromosome spreads from testes were prepared essentially as described (Alavattam et al. 2018). Testes were excised, detunicated, and placed in 1× phosphate-buffered saline (PBS). Seminiferous tubules were dissociated from whole testes. Seminiferous tubules were transferred to one well of a four-well dish (Thermo Fisher Nunc, 055047) containing 1 ml of cold PBS kept on ice. Seminiferous tubules were then gently unraveled into small clumps with fine-point tweezers; care was taken not to tear or mince the tubules. The clumps of seminiferous tubules were subsequently transferred to the second and third wells of 1 ml PBS for additional unraveling before transfer to the fourth well, which contained 1 ml hypotonic extraction buffer [HEB: 30 mM Tris base, 17 mM trisodium citrate, 5 mM ethylenediaminetetraacetic acid (EDTA), 50 mM sucrose, 5 mM dithiothreitol (DTT), and 1× cOmplete Protease Inhibitor Cocktail (Sigma, 11836145001). Once there, fine-point tweezers were used to carefully expose the surface area of tubules to HEB. The seminiferous tubules were incubated in HEB on ice for

30 min with gentle stirring every 10 min. After incubation, seminiferous tubules were mashed using a disposable surgical blade in 60 µL of 100 mM sucrose on a plain, uncharged microscope slide (Gold Seal: Thermo Fisher Scientific, 3010–002). After approximately 15–25 mashes, a semi-translucent cell suspension was formed. An additional 120 µL of sucrose (100 mM) was added to the suspension, and the suspension was mixed via gentle pipetting up and down several times. The diluted cell suspension was applied to positively charged slides (Probe On Plus: Thermo Fisher Scientific, 22-230-900) in volumes of 30 µL; before application of the suspension, the slides had been incubated in chilled fixation solution (2% paraformaldehyde, 0.1% Triton X-100, and 0.02% sodium monododecyl sulfate, adjusted to pH 9.2 with sodium borate buffer) for a minimum of 2 min. After applying the cell suspension/sucrose mixture, the slide was slowly and gently tilted up and down at slight angles (<10°) to mix the cell suspension/sucrose mixture with the remaining fixation solution. The slides were placed in "humid chambers" (closed pipet tip boxes filled to approximately two-thirds volume with water) at RT for a minimum of 1 h to maximum of overnight. Then, the slides were washed in a low-concentration surfactant, 0.4% Photo-Flo 200 (Kodak, 146– 4510), at RT two times, 2 min per wash. Slides were dried completely at RT (~30 min) before staining or storage at −80 °C.

**Immunostaining of meiotic chromosome spreads**

For immunostaining experiments, chromosome spreads were incubated in PBS containing 0.1% Tween 20 (PBST) for 5–30 min before blocking in antibody dilution buffer (PBST containing 0.15% BSA) for an additional 30–60 min. Primary and secondary antibodies (described below) were diluted in an antibody dilution buffer. Then, chromosome spreads were coated with 100 µL of the antibody/antibody dilution buffer solution, gently covered with Parafilm (Parafilm M All-Purpose Laboratory Film, Bemis Company, Inc.), and stored in humid chambers at 4 °C for a minimum period of 6 h to a maximum timespan of overnight (~15 h). This study made use of the following primary antibodies at the following dilutions [format: host anti-protein (source or company with product/catalog number if applicable), dilution]: rabbit anti-ATF7IP2 (generated for this study and recognizing amino acids 201–452 of mouse ATF7IP2), 1/500; mouse anti-SYCP3 (Abcam, ab97672), 1/5000; rabbit anti-SYCP3 (Novus, NB300–232), 1/500; rabbit anti-SYCP1 (Abcam, ab15090), 1/500; rabbit anti-H3K9me3 (Abcam, ab8898), 1/500; rabbit anti-H3K9me3 (Millipore, 07-442), 1/200; mouse anti-H3K9me2 (Abcam, ab1220), 1/500; rabbit anti-H3K9me1 (Millipore, 07-450), 1/500; rabbit anti-H3K9ac (Millipore, 06-942), 1/500; mouse anti-SYCP3 conjugated with Alexa-488 (Abcam, ab205846), 1/2000; rabbit anti-SCML2 (generated in the Namekawa Lab (Hasegawa et al. 2015)), 1/500; rabbit anti-SETDB1 (Proteintech, 11231-1-AP), 1/200; guinea pig anti-H1T (a gift from Dr. Mary Ann Handel (Inselman et al. 2003)), 1/500; rabbit anti-MLH1 (Santa Cruz, sc-11442), 1/200; mouse anti-TRIM28 (Active Motif, 61173), 1/250; rabbit anti-CHD4 (Active Motif, 39290), 1/100; mouse anti-H2AX-pS139 (γH2AX: Millipore, 05–636-1), 1/5000; mouse anti-H2AX-pS139 conjugated Alexa 647 (γH2AX: Millipore, 05–636-AF647), 1/5000; rabbit anti-ATF7IP (Abcam, ab84497), 1/500; sheep anti-MDC1 (ABD Sertec, AHP799), 1/500). After incubation with primary antibodies, slides were washed three times in PBST, 5 min per wash. Then, the slides were incubated with the appropriate secondary antibodies conjugated to Alexa 488, 555, or 647 fluorophores (Thermo Fisher Scientific). All secondary antibodies were diluted 1/500 in antibody dilution buffer. Slides were coated with 100 µL of the antibody/antibody dilution buffer solution; then, they were gently covered with Parafilm for 1 h incubation at RT in humid chambers in darkness. After slides were washed three times in PBST in darkness, 5 min per wash, they were counterstained with the DNA-binding chemical 4′,6-diamidino-2-phenylindole (DAPI; Sigma, D9542–5MG) diluted to 1 μg/ml concentration in PBS. Finally, slides were mounted using 20 μL undiluted ProLong Gold Antifade Mountant (Thermo Fisher Scientific, P36930). Slides were either imaged immediately or stored at 4 °C in darkness. Images were obtained with an ECLIPSE Ti-2 microscope (Nikon) equipped with an ORCA-Fusion sCMOS camera (Hamamatsu) and a 60× CFI Apochromat TIRF oil immersion objective NA 1.4 (Nikon), and were processed using NIS-Elements Basic Research (Nikon), Photoshop (Adobe), and Illustrator (Adobe) software.

**Judgement of pachytene substages**

Histone variant H1T is expressed starting in the mid-pachytene stage of male mouse meiosis (Drabent et al. 1996), and the duration of the pachytene stage is approximately 160 h (7 days) in mouse spermatogenesis (van der Heijden et al. 2007). H1T-negative early pachytene spermatocytes and H1T-positive mid-late pachytene spermatocytes were analyzed separately based on H1T immunostaining intensity. In addition to assessing H1T, we assessed pachytene substages by examining the morphology of chromosome axes stained with an anti-SYCP3 antibody as previously described (Alavattam et al. 2018).

**Histology and Immunostaining**

For the preparation of testis paraffin blocks, testes were fixed with 4% paraformaldehyde and 0.1% Triton X-100 in PBS at 4 °C overnight. Testes were dehydrated and embedded in paraffin. For histological analyses, 6 μm-thick paraffin sections were deparaffinized and stained with hematoxylin and eosin. For immunostaining, sections were autoclaved in Target Retrieval Solution, Citrate pH 6.1 (DAKO, S-1700) at 121 °C and 100 kPa (15 psi) for 10 min. The sections were blocked with Blocking One Histo (Nacalai USA, 06349–64) at RT for 30 min; the sections were then incubated with primary antibodies diluted in PBST at 4 °C overnight. The following antibodies were used at the following dilutions [format: host anti-protein (source or company with product/catalog number), dilution]: mouse anti-H2AX-pS139 (γH2AX: Millipore, 05–636), 1/5000; guinea pig anti-H1T (a gift from Dr. Mary Ann Handel (Inselman et al. 2003)), 1/500; mouse anti-TRIM28 (Abcam, ab109287), 1/500; rabbit anti-ATF7IP (Abcam, ab84497), 1/500; rabbit anti-STRA8 (Abcam, ab49602), 1/500; goat anti-ZBTB16/PLZF (R&D Systems, AF2944), 1/500; goat anti-CD117/c-KIT (R&D Systems, AF1356), 1/500; mouse anti-H2AX-pS139 (γH2AX: Millipore, 05–636), 1/5000; mouse anti-H2AX-pS139 conjugated Alexa 647 (γH2AX: Millipore, 05–636-AF647), 1/5000. The resulting signals were detected with secondary antibodies conjugated to Alexa 488 or 555 (Thermo Fisher Scientific), diluted 1/1000 in PBST, and incubated at RT for 1 h. Sections were counterstained with DAPI as described above. Images were obtained with a Zeiss LSM800 confocal microscope equipped with an Axiocam 506 monochrome camera (Zeiss) and a C-Apochromat 40x/1.2 water lens (Zeiss), and were processed using Fiji (NIH) (Schindelin et al. 2012), Photoshop (Adobe), and Illustrator (Adobe) software.

**Isolation of pachytene spermatocytes**

Isolation of pachytene spermatocytes using Fluorescence-activated cell sorting (FACS) was performed using SH800S (SONY), with Vybrant DyeCycle Violet Stain (DCV) (Invitrogen, V35003) stained testicular single-cell suspensions prepared as described previously (Yeh et al. 2021). For H3K9 CUT&RUN experiments and bulk RNA-seq experiments, early-to-mid pachytene spermatocytes were isolated from P15 *Atf7ip2*$^{+/+}$ mice, when germ cells reach early-mid pachynema in the first wave of spermatogenesis. As spermatocytes are arrested at mid-pachynema in *Atf7ip2*$^{-/-}$ mice, >P15 mice were used for isolating arrested pachytene spermatocytes from *Atf7ip2*$^{-/-}$ mice. For ATF7IP2 CUT&Tag experiments and H3K9 CUT&RUN experiments using wild-type pachytene spermatocytes, adult C57BL/6J mice were used for isolation of pachytene spermatocytes.

In brief, to prepare single cells in suspension for cell sorting, detangled seminiferous tubules from adult mouse testes were (1) washed with 1× Hanks' Balanced Salt Solution (HBSS, Gibco, 14175095) three times, (2) incubated with 1 mg/ml Type 1 collagenase (Worthington, CLS1) for 20 min at 35 °C to remove interstitial cells, and (3) digested in Dulbecco's Modified Eagle Medium (DMEM, Gibco, 11885076) containing 2% Fetal Bovine Serum (FBS, Gibco, 16000044), 2 mg/ml Type 1 collagenase, 0.25 mg/ml DNase I (Sigma, D5025), 1.5 mg/ml Hyaluronidase (Sigma, H3506) and 700 U/ml Recombinant Collagenase (FUJIFILM Wako Pure Chemical Corporation, 036-23141) at 35 ℃ for 20–30 min with vigorous pipetting for cell dissociation. The resulting cell suspension was plated in Lectin from Datura stramonium (Sigma L2766)-coated plates for 20 min in DMEM/F-12 medium (Gibco, 11330107) supplemented with 10% fetal bovine serum (FBS), which promoted the adhesion of the remaining somatic cells. The floating germ cell-enriched cell suspension was then collected from plates and washed with 10 ml FACS buffer (PBS containing 2% FBS) three times by centrifugation at 300 × g for 5 min and filtered through a 70 µm nylon cell strainer (Falcon, 352350). The resultant single cells were stained with DCV. After 30 min incubation at 35 ℃, cells were filtered into a 5 ml FACS tube through a 35 µm nylon mesh cap (Falcon, 352235). Samples were kept on ice until sorting.

**Single-cell RNA-seq library generation**

Testes were collected from P15 *Atf7ip2*$^{+/+}$ and *Atf7ip2*$^{-/-}$ mice. In brief, to prepare a suspension of individual testicular cells, detangled seminiferous tubules from adult mouse testes were first washed with 1× Hanks' Balanced Salt Solution (HBSS, Gibco, 14175095) three times and then incubated in Dulbecco's Modified Eagle Medium (DMEM, Gibco, 11885076) containing 2% Fetal Bovine Serum (FBS, Gibco, 16000044), 2 mg/ml Type 1 collagenase (Worthington, CLS1), 0.25 mg/ml DNase I (Sigma, D5025), 1.5 mg/ml Hyaluronidase (Sigma, H3506) and 700 U/ml Recombinant Collagenase (FUJIFILM Wako Pure Chemical Corporation, 036-23141) at 35 ℃ for 20 min, then dissociated using vigorous pipetting and incubated at 35 ℃ for another 10 min. The single cell suspension was then washed with 10 ml FACS buffer (PBS containing 2% FBS) three times by centrifugation at 300 × g for 5 min and filtered into a 5 ml FACS tube through a 35 µm nylon mesh cap (Falcon, 352235). To exclude dead cells, 7-AAD Viability Stain (Invitrogen, 00-6993-50) was added to the cell suspension. After removing 7-AAD+ dead cells, cells were collected using a Cell Sorter SH800 (SONY).

Collected cells were resuspended in 1× PBS containing 2% BSA and submitted to the DNA Technologies & Expression Analysis Core of the University of California, Davis, for library preparation. Briefly, approximately 10,000 single cells were loaded on the Chromium Controller (10X Genomics Inc.). Single-cell RNA-seq libraries were generated using Chromium Single Cell 3' Reagent Kits (v3) following the manufacturer's instructions and sequenced as 75-bp paired-end reads with an Element Bio AVITI system.

**Bulk RNA-seq library generation with ERCC RNA spike-in for isolated pachytene spermatocytes**

*Atf7ip2*$^{+/+}$ and *Atf7ip2*$^{-/-}$ pachytene spermatocyte RNA-seq libraries were prepared using the Smart-seq2 method (Picelli et al. 2014) as follows: ~10,000 isolated cells were used as one replicate. Two independent biological replicates were used per RNA-seq library generation. External RNA Controls Consortium (ERCC) RNA Spike-In Mix (Invitrogen, 4456740) was added to cell lysis before RNA extraction; first, ERCC RNA Spike-In Mix was diluted 1000 times with nuclease-free water, and then 1 µl was added to 10,000 cells.

Total RNA was extracted using the RNeasy Plus Micro Kit (QIAGEN, Cat #74034) according to the manufacturer's instructions. Libraries were constructed using a NEBNext® Single Cell/Low Input RNA Library Prep Kit for Illumina® (NEB, E6420S) according to the manufacturer's instructions. Prepared RNA-seq libraries were sequenced as 150-bp paired-end reads with a Hiseq 4000 system (Illumina).

**Bulk RNA-seq library generation for juvenile testes samples**

For testis bulk RNA-seq library preparation, total RNA was purified from *Atf7ip2*$^{+/+}$ or *Atf7ip2*$^{-/-}$ whole testis using an RNeasy Plus Micro Kit (QIAGEN, Cat # 74034) according to the manual provided. Two independent biological replicates were used for RNA-seq library generation. RNA quality and quantity were checked using, respectively, a Bioanalyzer (Agilent) and Qubit (Life Technologies). Library construction was performed by the CCHMC DNA Sequencing and Genotyping Core (Cincinnati, Ohio, USA) using the Illumina TruSeq stranded mRNA kit after polyA enrichment. Prepared RNA-seq libraries were sequenced as 100-bp paired-end reads with a Novaseq 6000 system (Illumina).

**CUT&RUN/Tag library generation**

*Atf7ip2*$^{+/+}$ and *Atf7ip2*$^{-/-}$ H3K9me3 pachytene spermatocyte CUT&RUN libraries were generated as previously described (Skene et al. 2018). A step-by-step protocol is available at protocols.io: dx.doi.org/10.17504/protocols.io.zcpf2vn. Approximately 10,000 FACS-sorted pachytene spermatocytes were used as one replicate, and two independent biological replicates were used. The antibody used was rabbit anti-H3K9me3 (Abcam; ab8898), 1/100. Homemade Protein A/G-MNase was provided by the laboratory of Dr. Artem Barski. Spike-in *E. coli* DNA was included with each reaction by adding 0.4 ng per 10,000 cells. Libraries were constructed using a NEBNext Ultra II DNA Library Prep Kit for Illumina (NEB, E7645S). Prepared CUT&RUN libraries were sequenced as 150-bp paired-end reads on a Hiseq 4000 system (Illumina).

Wild-type ATF7IP2 (rabbit anti-ATF7IP2, 1/100) pachytene spermatocyte CUT&Tag libraries were prepared as previously described (Kaya-Okur et al. 2019; Kaya-Okur et al. 2020) using CUTANA™ pAG-Tn5 (Epicypher, 15-1017). A step-by-step protocol is available at protocols.io: dx.doi.org/10.17504/protocols.io.bcuhiwt6. CUT&Tag libraries were sequenced as 150-bp paired-end reads on a HiSeq 4000 system (Illumina).

**ScRNA-seq data processing**

Fastq files were processed and aligned to the mouse mm10 transcriptome (GENCODE vM23/Ensembl 98) using the 10X Genomics Cell Ranger v 7.0.0 pipeline. Further analyses were conducted with R (ver.3.6.2) via RStudio (ver.1.2.1335). Quality assessment of scRNA-seq data and primary analyses were conducted using the Seurat package for R (v3.2.2) (Butler et al. 2018) (Stuart et al. 2019). To remove the effect of low-quality cells, only cells that expressed more than 200 genes were used for further analyses. The scRNA-seq data were merged and normalized using the Seurat function SCTransform. Using the function RunUMAP, both dimensionality reduction analysis and cluster visualization were conducted. The clustering of cells was conducted using the functions FindNeighbors and FindClusters with default settings. The function FindMarker was called with default settings to characterize groups of cells. For the identification of marker genes in each cluster and to determine differentially expressed genes (DEGs), the FindAllMarkers function was used with the parameters only.pos = T, min.pict = 0.25, and logfc.threshold = 0.25. To extract the germ-cell population, only cell populations expressing the germ cell marker *Ddx4* were selected. The selected cell populations were plotted in a UMAP representation. The cell populations showing the expression of somatic cell marker genes and the enrichment of mitochondrial genes were removed, and the remaining cells were used as germ-cell populations. RNA velocity analysis was conducted with default settings using the Python package velocity package (Python 3.7.3) and R (La Manno et al. 2018), and visualized in UMAP representations.

Gene enrichment analyses were performed using Metascape (Zhou et al. 2019) with default settings, and the results were visualized using R and GraphPad Prism9. To characterize each cluster, the top 100 representative genes per cluster were used.

Genes located on autosomes and sex chromosomes were obtained from an mm10 GTF file. Averaged autosomal and sex chromosomal gene expression values were calculated using per-gene normalized values.

**Bulk RNA-seq with ERCC spike-in data processing**

Raw RNA-seq reads were trimmed with the program Trim Galore (version 0.6.6; called with flag --paired) and then aligned to the mouse (GRCm38/mm10) genome using STAR (version 2.5.4b) (Dobin et al. 2013). Using featureCounts (Liao et al. 2014) (version 2.0.1), raw counts matrices were generated with respect to a concatenated UCSC mouse RefSeq and ERCC annotation. ERCC spike-in normalization was performed with the packages edgeR (Robinson et al. 2010) and Limma (Ritchie et al. 2015). First, normalization factors were calculated using the number of reads mapped to ERCC sequences. Filtered raw reads (>2 in at least one sample) were assessed for library size using normalization factors and then voom-transformed. The topTable function was used to identify differentially expressed genes (DEGs: Benjamini-Hochberg-adjusted p-value $< 0.05$; log2 fold change $> 2$). To perform GO analyses of DEGs, we used the online functional annotation clustering tool Metascape (Zhou et al. 2019).

**Testis bulk RNA-seq data processing**

Raw RNA-seq reads were trimmed and filtered using trimmomatic (version 0.39) (Bolger et al. 2014) and then aligned to the mouse genome (GRCm38/mm10) using STAR (version 2.5.4b) (Dobin et al. 2013) with parameter --outFilterMultimapNmax 1 to contain only unique alignments. For analyses, the sequencing data were uploaded to the Galaxy web platform (Afgan et al. 2016). To identify DEGs between control and experimental conditions, raw read count files were generated using featureCounts (Liao et al. 2014) (version 2.0.1) based on the UCSC mouse RefSeq annotation. After quantification, a counts matrix was input to DESeq2 (Love et al. 2014) (version. 1.38.2) for differential gene expression analyses. Differentially expressed genes were identified with cutoffs ≥ 1.5-fold change and Benjamini-Hochberg-adjusted p-values of ≤ 0.05 among expressed genes (TPM > 1 in either the *Atf7ip2*$^{+/+}$ or the *Atf7ip2*$^{-/-}$). P-values for autosome and sex chromosome comparisons were obtained from hypergeometric tests. Average fold changes for autosomes and sex chromosomes were calculated from the per-chromosome TPM averages for *Atf7ip2*$^{+/+}$ or *Atf7ip2*$^{-/-}$. Genes with TPM > 1 in the *Atf7ip2*$^{+/+}$ or the *Atf7ip2*$^{-/-}$ were defined as expressed genes and used in analyses. Plots of differentially expressed genes for autosomes and sex chromosomes were produced using ggplot2 (3.4.0) in R. For gene ontology analyses, the Database for Annotation, Visualization and Integrated Discovery (DAVID) (v2022q3) was used to identify the GO terms (Huang da et al. 2009; Sherman et al. 2022). Heatmaps of expression profiles were produced using Morpheus (https://software.broadinstitute.org/morpheus).

**CUT&RUN/Tag data processing**

For CUT&RUN/Tag data processing, the steps in a data analysis tutorial (dx.doi.org/10.17504/protocols.io.bjk2kkye) were followed. Briefly, after trimming with the program Trim Galore (version 0.6.6), reads were aligned to the mouse genome (GRCm38/mm10) using Bowtie2 (Langmead and Salzberg 2012) (version 2.4.2) with options: --end-to-end --very-sensitive --no-mixed --no-discordant --phred33 -I 10 -X 700. For mapping *E.coli* spike-in fragments, the --no-overlap --no-dovetail options were called to avoid cross-mapping. PCR duplicates were removed using the Picard Tools MarkDuplicates command (version 2.23.8) (https://broadinstitute.github.io/picard/). To compare replicates, Pearson correlation coefficients were calculated and plotted with the deepTools commands multiBamSummary and plotCorrelation from deepTools (Ramirez et al. 2016) (version 3.5.0). Spike-in normalization was implemented using the exogenous scaling factor computed from the *E.coli* mapping files, where scaling factors were equal to 1,000,000 ÷ number of spike-in alignments.

To call "peaks" of local CUT&Tag ATF7IP2 enrichment, SEACR (Meers et al. 2019) (https://seacr.fredhutch.org/) was used. HOMER (Heinz et al. 2010) (version 4.11) was used for peak annotation. Genes with promoter-associated ATF7IP2 peaks were identified as ATF7IP2 target genes.

For visualization using the Integrative Genomics Viewer (Robinson et al. 2011) (version 2.5.3), spike-in-normalized genome coverage tracks for *Atf7ip2*$^{+/+}$ and *Atf7ip2*$^{-/-}$ pachytene spermatocyte H3K9me3 data were generated using the bedtools (Quinlan and Hall 2010) (version 2.29.2) function genomecov with options -bg -scale ${scale_factor}. The program deepTools (Ramirez et al. 2016) was used to draw tag density plots and heatmaps.

**Analysis of TE expression**

For the evaluation of TE expression, we used the aligner STAR to align RNA-seq reads to the mouse (GRCm38/mm10) genome allowing only uniquely mapped reads. Using the program featureCounts, raw counts matrices were generated with respect to a custom repetitive-sequence annotation termed a "best-match" TE annotation (Sakashita et al. 2020), the use of which enables the detection of alignments uniquely mapped to TEs that are not exon-derived (mRNA-derived) and thus excludes TEs that overlap genes. The use of this best-match TE annotation mitigates the potential for erroneously counting intron-derived reads as TEs. Filtered raw reads (>2 in at least one sample) were run through DESeq2 (Love et al. 2014) differential expression analyses ( Benjamini-Hochberg-adjusted p-value < 0.05; log2 fold change > 2). To perform spike-in normalization, the function estimateSizeFactor was run with argument controlGenes set to the ERCC features.

**Quantification and statistical analyses**

No data were excluded from analyses. The experiments were not randomized, and investigators were not blinded to allocation during experiments and outcome assessment. Where appropriate, the mean is reported as a measurement of central tendency, and the SEM is used as a measure of precision. Statistical significance is threshold at $\alpha = 0.05$; p-values $< \alpha$ are considered significant. Details for statistical analyses performed in this study are described in relevant portions of the Results section and figure legends. Sample sizes used for analyses are described in relevant portions of the Results section and figure legends. In predetermining sample sizes, we sought to analyze a minimum of three independent samples; the only exception is bulk and single-cell RNA-seq experiments, and CUT&RUN/Tag experiments, all which were performed with two independent samples. Measurements were recorded in Excel (Microsoft) and R.

**Code availability**

Source code for all software and tools used in this study—with documentation, examples, and additional information—is available at the URLs listed below.

trimmomatic [http://www.usadellab.org/cms/?page=trimmomatic]
STAR [https://github.com/alexdobin/STAR]
bowtie2 [https://github.com/BenLangmead/bowtie2]
bedtools[https://github.com/arq5x/bedtools2]
featureCounts [http://subread.sourceforge.net]
DESeq2 [https://bioconductor.org/packages/release/bioc/html/DESeq2.html]
ggplot2 [https://github.com/tidyverse/ggplot2]
deepTools [https://deeptools.readthedocs.io/en/develop/index.html]
Morpheus [https://software.broadinstitute.org/morpheus]
DAVID [https://david.ncifcrf.gov]
Galaxy [https://usegalaxy.org]
Hypergeometric p-value calculator [https://systems.crump.ucla.edu/hypergeometric/]

**A** Expresson of *Atf7ip2* in mouse tissues

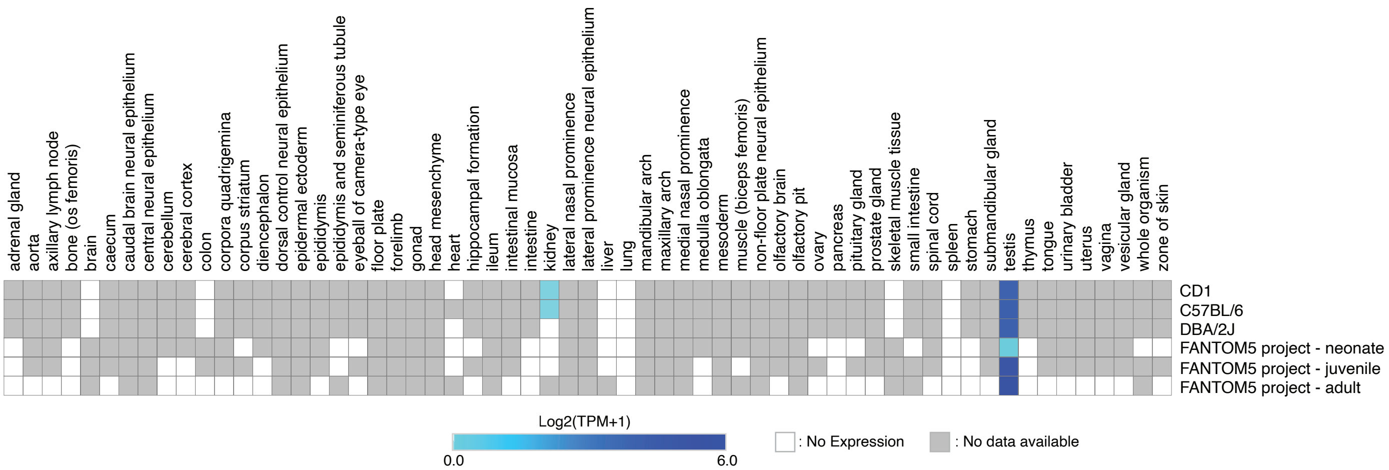


**B** Amino acid sequences of human ATF7IP2 (682 aa) and mouse ATF7IP2 (452 aa)

```
Human      MASPDRSKRKILKAKKTMPLSCRKQVEMLNKSRNVEALKTAIGSNVPSGNQSFSPSVITR      60
Mouse      ------------------------------------------------------------

Human      TTEITKCSPSENGASSLDSNKNSISEKSKVFSQNCIKPVEEIVHSETKLEQVVCSYQKPS      120
Mouse      ------------------------------------------------------------      0

Human      RTTESPSRVFTEEAKDSLNTSENDSEHQTNVTRSLFEHEGACSLKSSCCPPSVLSGVVQM      180
Mouse      ------------------------------------------------------------      0

Human      PESTVTSTVGDKKTDQMVFHLETNSNSESHDKRQSDNILCSEDSGFVPVEKTPNLVNSVT      240
Mouse      --------------------------MESPDRKR-QKVLKAKK--TMPTS----------      21
                                       ** *::: :::* ::.   :*..

Human      SNNCADDILKTDECSRTSISNCESADSTWQSSLDTNNNSHYQKKRMFSENEENVKRMKTS      300
Mouse      -YQKQLEIILNK-------STNVEAPKTT-----VGTNIPNGHNQKMFSKNKENVKVMKVS      68
             :   :**:.         :* *: .:*        .*  : ::::***:*:**** **.*

Human      EQINENICVSLERQTAFLEQVRHLIQQEIYSINYELFDKKLKELNQRIGKTECRNKHEGI      360
Mouse      EQINENACGALERHTALLEQVKHWIRQEICMINCNLFDKKLNELNERIGKTQCKSRHEAI      128
           ****** * :***:**:****:* *:***   ** :******:***:*****:*:.:**.*

Human      ADKLLAKIAKLQRRIKTVLLFQRNCLKPNMLSSNGASKVANSEAMILDKNLESVNSPI-E      419
Mouse      AGELFVKIRRLQKRIKTVLSSQRNCLEPNTLPSNTVCKVTDSEAMNLNVTQKSVKSRSKR      188
           *.:*:.** :**:******  *****:** * ** ..**::**** *: . :**:*   .

Human      KSSVNYEPSNPSEKGSKKINLSSDQNKSVSESNNDDVMLISVESPNLTTPITSNPTDTRK      479
Mouse      ISSVNHTPLNSSEKAGRKTNLPSTCVEFASESNTDDVMLISVKNSNLTTSITSEQTEIRK      248
            ****: * * ***..:* ** *    : .****.********:. **** ***: *: **

Human      ITSGNSSNSPNAE--VMAVQKKLDSIIDLTKEGLSNCNTESPVSPLESHSKAASNSKETT      537
Mouse      NTSRNLSNSPNSMIKVGPVEKKFDFVIDLTREGPSNYSIESPSFTLKSTSKAVLRSKEII      308
            ** * *****:    *   *:**:* :****:** ** . ***    *:* ***. .***

Human      PLAQNAVQVPESFEHLPPLPEPPAPLPELVDKTRDTLPPQKPELKVKRVFRPNGIALTWN      597
Mouse      PVAENGNEGFGSFEHLPPLPEPPAPLPEMADKIKDTLPPQKPELKVKWVLRPTSIALTWN      368
           *:*:*. :    ****************:.** :************* *:**..******

Human      ITKINPKCAPVESYHLFLCHENSNNKLIWKKIGEIKALPLPMACTLSQFLASNRYYFTVQ      657
Mouse      IPKVNPNCAPVESYHLFLYYENS-DHLTWKKIAEIKALPLPMACTLSQNLASTKYYFAVQ      427
           * *:**:*********** :*** ::* ****.************** ***.:***:**

Human      SKDIFGRYGPFCDIKSIPGFSENLT       682
Mouse      SKDIFGRYGPFCNIKSIPRFSENLT       452
```

**C** Human testicular sections

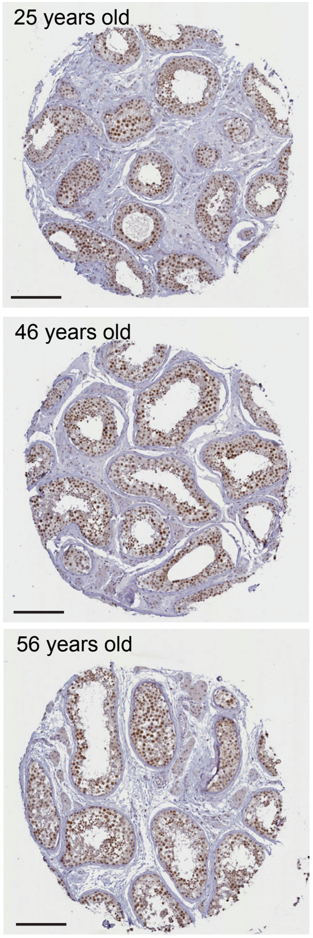


ATF7IP2: DAB staining

**Supplemental Figure S1. ATF7IP2 expression patterns in mouse tissues and human testes.**
**(A)** *Atf7ip2* expression levels in mouse tissues. Data were sourced and adapted from the Expression Atlas (https://ww-w.ebi.ac.uk/gxa/home).
**(B)** Human and mouse ATF7IP2 amino acid sequences via Clustal Omega alignment.
**(C)** Representative testis sections from humans at 25, 46, and 56 years of age immunohistochemically stained with an antibody raised against ATF7IP2 (brown) and counterstained with hematoxylin. Images of human testis sections were sourced and adapted from the Human Protein Atlas (https://www.proteinatlas.org/). Scale bars: 200 µm.

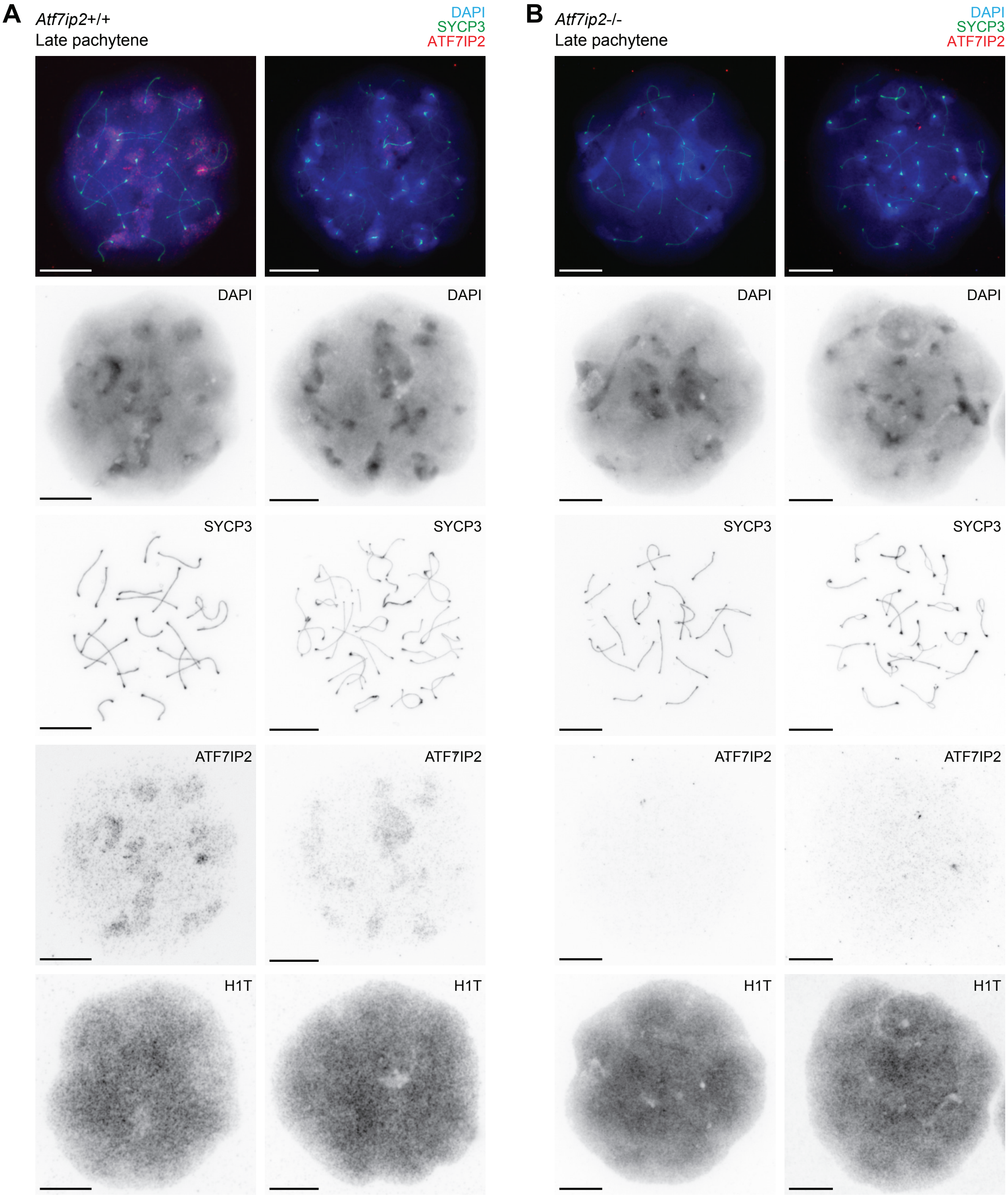


**Supplemental Figure S2. ATF7IP2 protein is absent from *Atf7ip2*$^{-/-}$ spermatocytes.**
**(A, B)** Chromosome spreads of *Atf7ip2*$^{+/+}$ **(A)** and *Atf7ip2*$^{-/-}$ **(B)** spermatocytes stained with DAPI and antibodies raised against SYCP3 (a marker of chromosome axes), ATF7IP2, and H1T (a marker of germ cells in mid pachytene and subsequent stages). Scale bars: 10 µm.

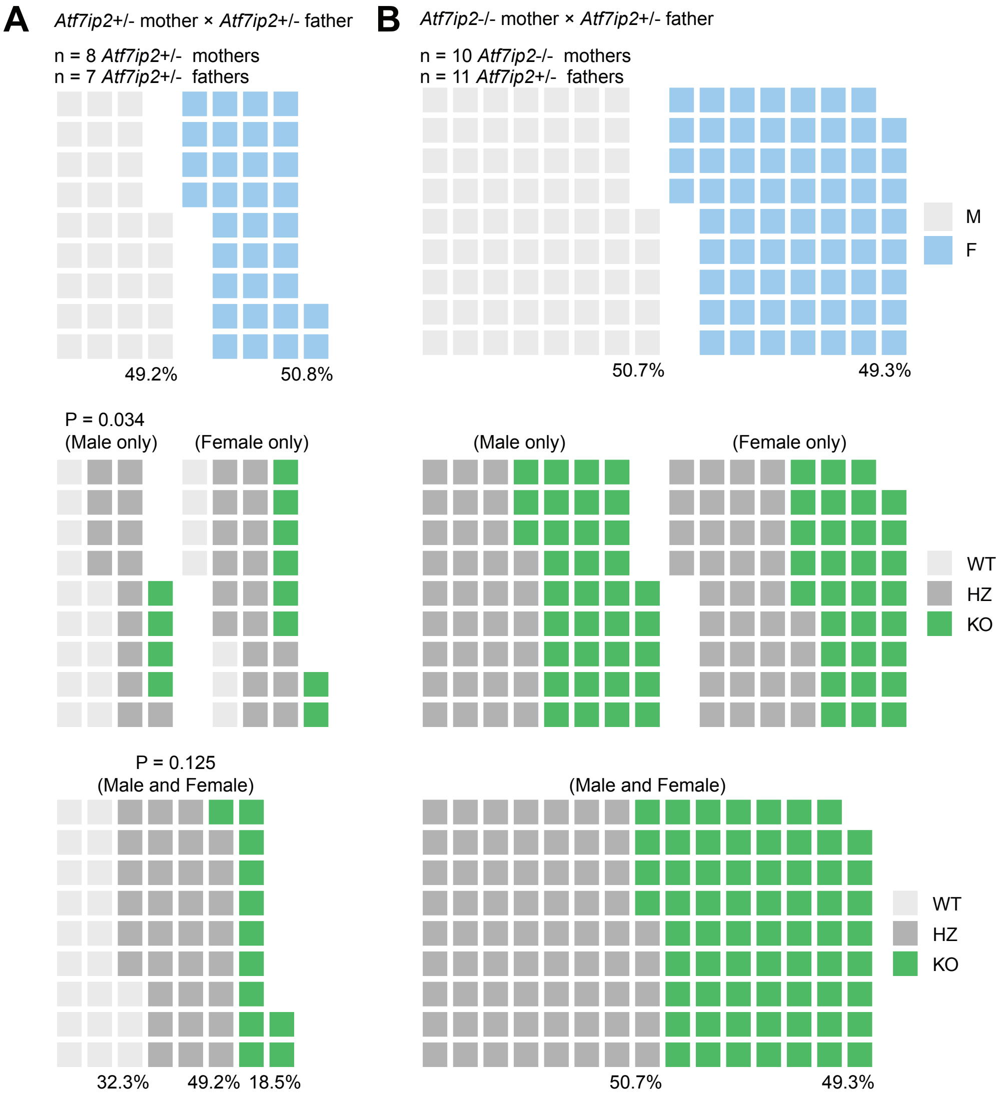


**Supplemental Figure S3. Breeding summary of *Atf7ip2* knockout mice.**
**(A, B)** Breeding summary of *Atf7ip2*$^{+/-}$ mothers and *Atf7ip2*$^{+/-}$ fathers **(A)**, and *Atf7ip2*$^{+/-}$ mothers and *Atf7ip2*$^{+/-}$ fathers **(B)**. Each square represents an individual offspring. Numbers of mice used for breeding are shown. P-values are from chi-squared tests.

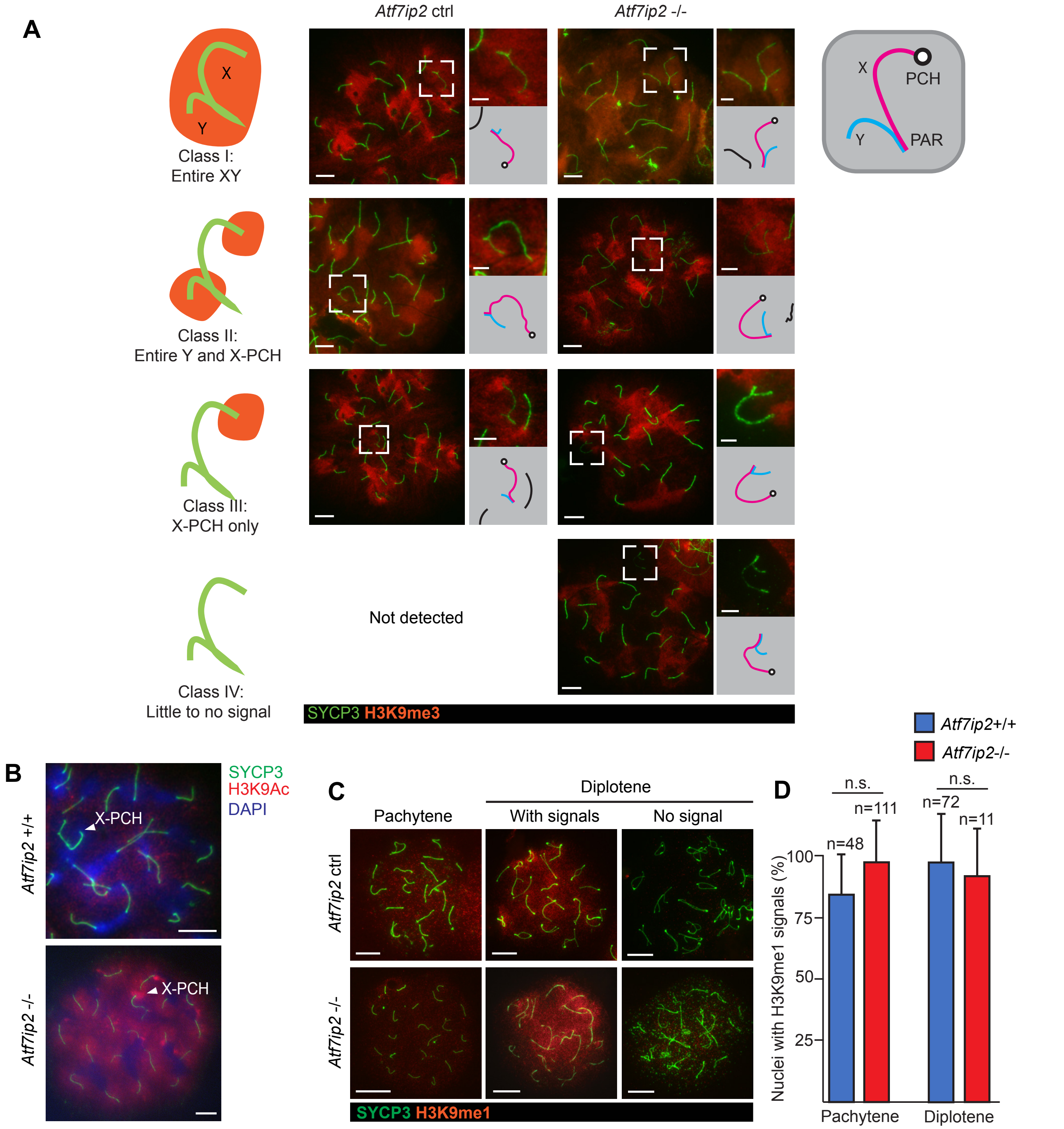


**Supplemental Figure S4. H3K9me3, H3K9ac, and H3K9me1 localization in *Atf7ip2*$^{-/-}$ spermatocytes.**

**(A)** H3K9me3 accumulation patterns on the sex chromosomes in early pachytene spermatocytes. H3K9me3 distribution patterns are categorized into four classes. The four classes are organized in order of decreasing H3K9me3 enrichment on the XY domain. Class I: broad, uninterrupted H3K9me3 accumulation on the XY chromosomes; Class II: enrichment of H3K9me3 on X pericentric heterochromatin (PCH) and chromosome Y; Class III: H3K9me3 enrichment on X-PCH only. Class IV: little-to-no H3K9me3 accumulation on XY. Chromosome spreads of *Atf7ip2* control (*Atf7ip2* ctrl: *Atf7ip2*$^{+/+}$ and *Atf7ip2*$^{+/-}$) and *Atf7ip2*$^{-/-}$ spermatocytes stained with antibodies raised against H3K9me3 and SYCP3 (a marker of chromosome axes). Dashed squares are magnified in the panels to the upper right, and XY schema are shown in the lower right. Scale bars: 10 µm for whole nuclei, 5 µm for XY.

**(B)** *Atf7ip2* Ctrl and *Atf7ip2*$^{-/-}$ spermatocyte chromosome spreads stained with antibodies raised against H3K9ac and SYCP3. Scale bars: 10 µm.

**(C)** *Atf7ip2* Ctrl and *Atf7ip2*$^{-/-}$ spermatocyte chromosome spreads stained with antibodies raised against H3K9me1 and SYCP3. Representative photos for nuclei with and without H3K9me1 signal are shown for diplotene spermatocytes. Scale bars: 10 µm.

**(D)** Quantification of pachytene and diplotene spermatocytes with H3K9me1 signals on the sex chromosomes. Three independent experiments. P-values are from Fisher's exact tests: n.s. (not significant).

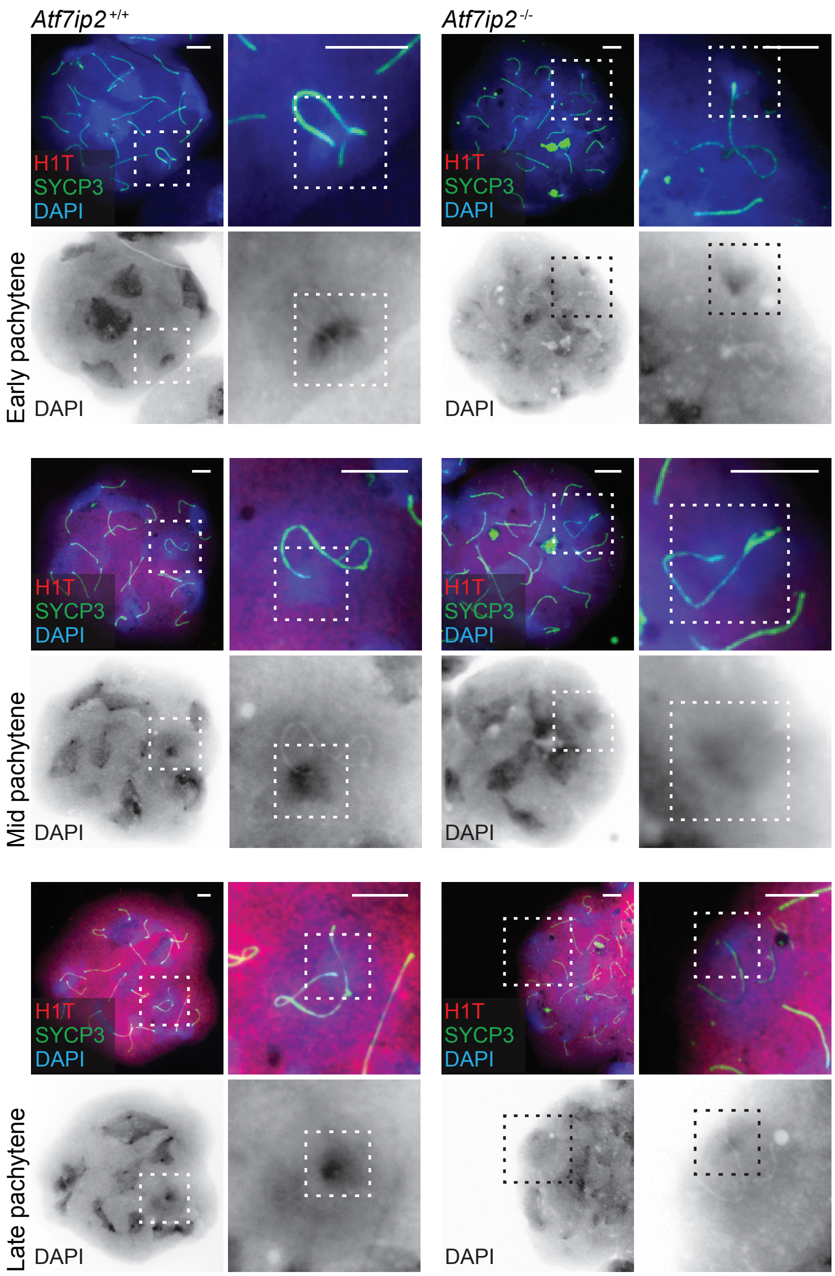


**Supplemental Figure S5. X-PCH defects in *Atf7ip2*$^{-/-}$ spermatocytes.**
Chromosome spreads of *Atf7ip2*$^{+/+}$ and *Atf7ip2*$^{-/-}$ spermatocytes stained with DAPI and antibodies raised against SYCP3 and H1T. Scale bars: 10 µm. Dashed squares are magnified in the panels to the right. In comparison to controls, *Atf7ip2*$^{-/-}$ X-PCH DAPI intensity is decreased.

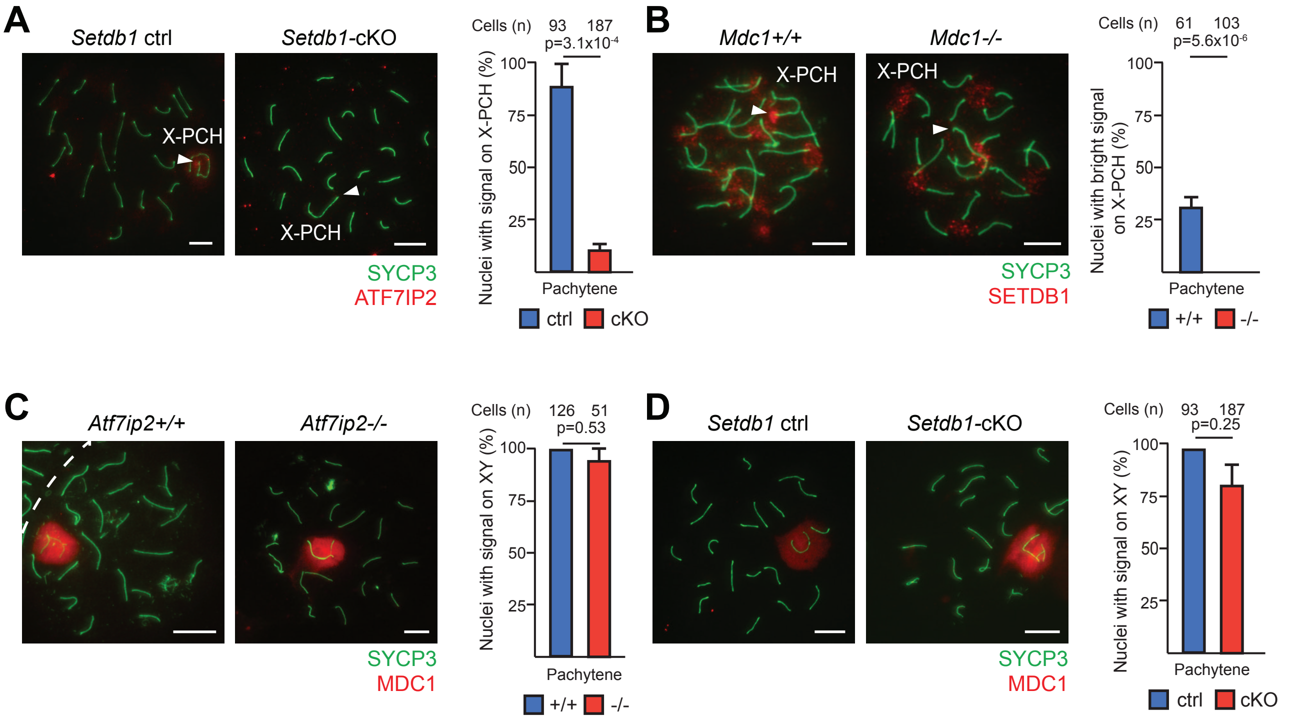


**Supplemental Figure S6. Evaluation of *Setdb1*-cKO, *Mdc1*$^{-/-}$, and *Atf7ip2*$^{-/-}$ spermatocytes**

**(A, D)** *Setdb1* ctrl and Setdb1-cKO spermatocyte chromosome spreads stained with DAPI and antibodies raised against SYCP3 and ATF7IP2 (A) and MDC1 (D).

**(B)** *Mdc1*$^{+/+}$ and *Mdc1*$^{-/-}$ spermatocyte chromosome spreads stained with DAPI and antibodies raised against SYCP3 and SETDB1.

**(C)** *Atf7ip2*$^{+/+}$ and *Atf7ip2*$^{-/-}$ spermatocyte chromosome spreads stained with DAPI and antibodies raised against SYCP3 and MDC1.

Scale bars: 10 µm.

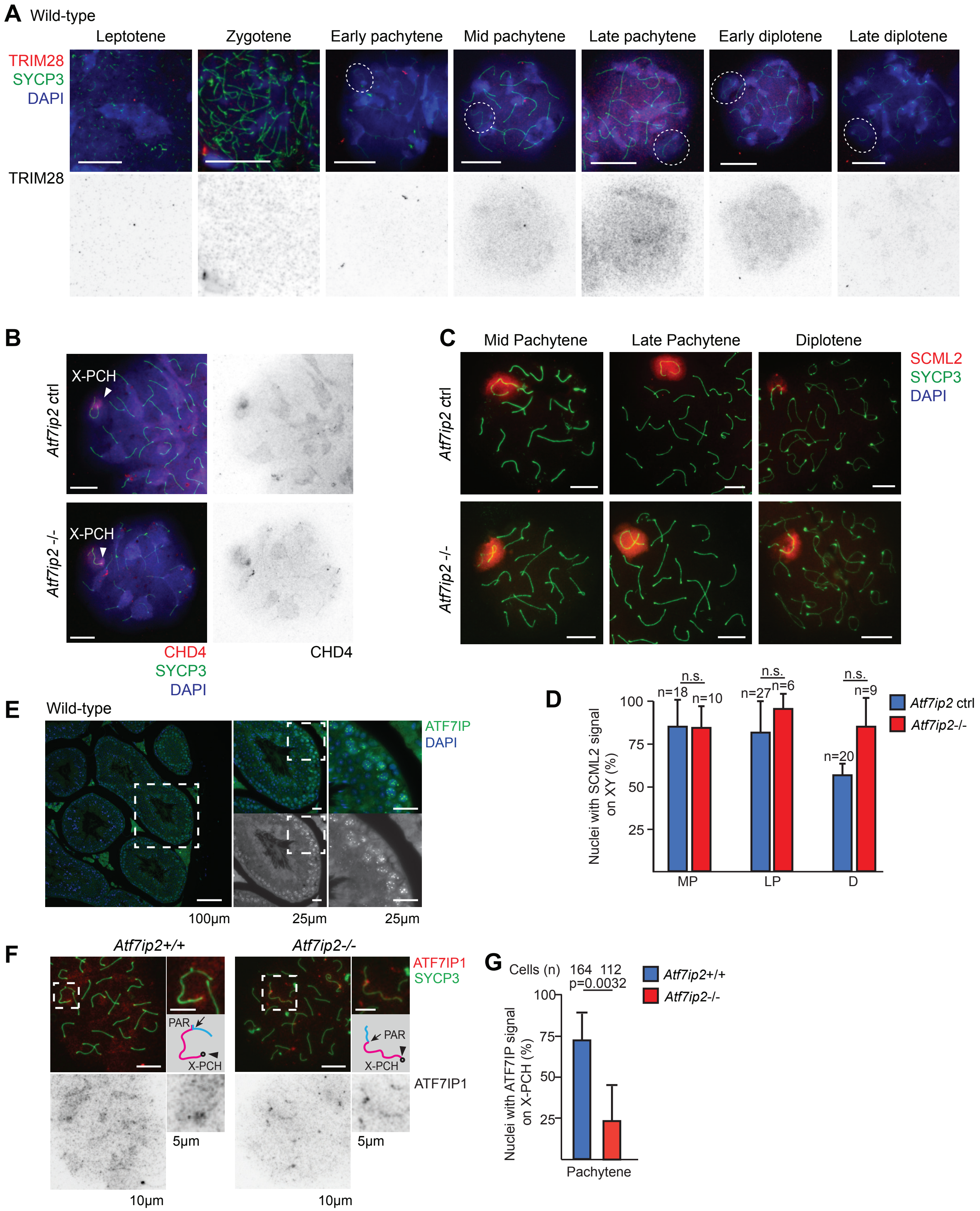

A Wild-type
Leptotene
Zygotene
Early pachytene
Mid pachytene
Late pachytene
Early diplotene
Late diplotene
TRIM28
SYCP3
DAPI
TRIM28
B
Atf7ip2 ctrl
X-PCH
Atf7ip2 -/-
X-PCH
CHD4
SYCP3
DAPI
CHD4
C
Mid Pachytene
Late Pachytene
Diplotene
SCML2
SYCP3
DAPI
Atf7ip2 ctrl
Atf7ip2 -/-
D
Nuclei with SCML2 signal on XY (%)
100
75
50
25
n.s.
n.s.
n.s.
n=18
n=10
n=27
n=6
n=20
n=9
MP
LP
D
Atf7ip2 ctrl
Atf7ip2-/-
E
Wild-type
ATF7IP
DAPI
100µm
25µm
25µm
F
Atf7ip2+/+
Atf7ip2-/-
ATF7IP1
SYCP3
PAR
X-PCH
PAR
X-PCH
ATF7IP1
5µm
5µm
10µm
10µm
G
Cells (n)
164
112
p=0.0032
Atf7ip2+/+
Atf7ip2-/-
Nuclei with ATF7IP signal on X-PCH (%)
100
75
50
25
Pachytene

**Supplemental Figure S7. TRIM28, CHD4, SCML2, and ATF7IP localization in chromosome spreads.**
**(A)** Wild-type spermatocyte chromosome spreads stained with antibodies raised against SYCP3 and TRIM28. Scale bars: 10 μm.
**(B)** *Atf7ip2* ctrl and *Atf7ip2*$^{-/-}$ spermatocyte chromosome spreads stained with antibodies raised against SYCP3 and CHD4. Scale bars: 10 μm.
**(C)** *Atf7ip2* ctrl and *Atf7ip2*$^{-/-}$ spermatocyte chromosome spreads stained with antibodies raised against SYCP3 and SCML2. Scale bars: 10 μm.
**(D)** Quantification of spermatocytes with SCML2 signals on the sex chromosomes. Three independent experiments. P-values are from Fisher's exact tests: n.s.: not significant.
**(E)** Wild-type testis sections from mice at 4 months of age stained with DAPI and antibodies raised against ATF7IP. Dashed squares are magnified in the panels to the right. Scale bars: 100 μm (left panel) and 25 μm (magnified panels).
**(F)** *Atf7ip2*$^{+/+}$ and *Atf7ip2*$^{-/-}$ spermatocyte chromosome spreads stained with antibodies raised against SYCP3 and ATF7IP. Scale bars: 10 μm.
**(G)** Quantification of spermatocytes with X-PCH ATF7IP signals. Three independent experiments. P-value is from Fisher's exact tests.

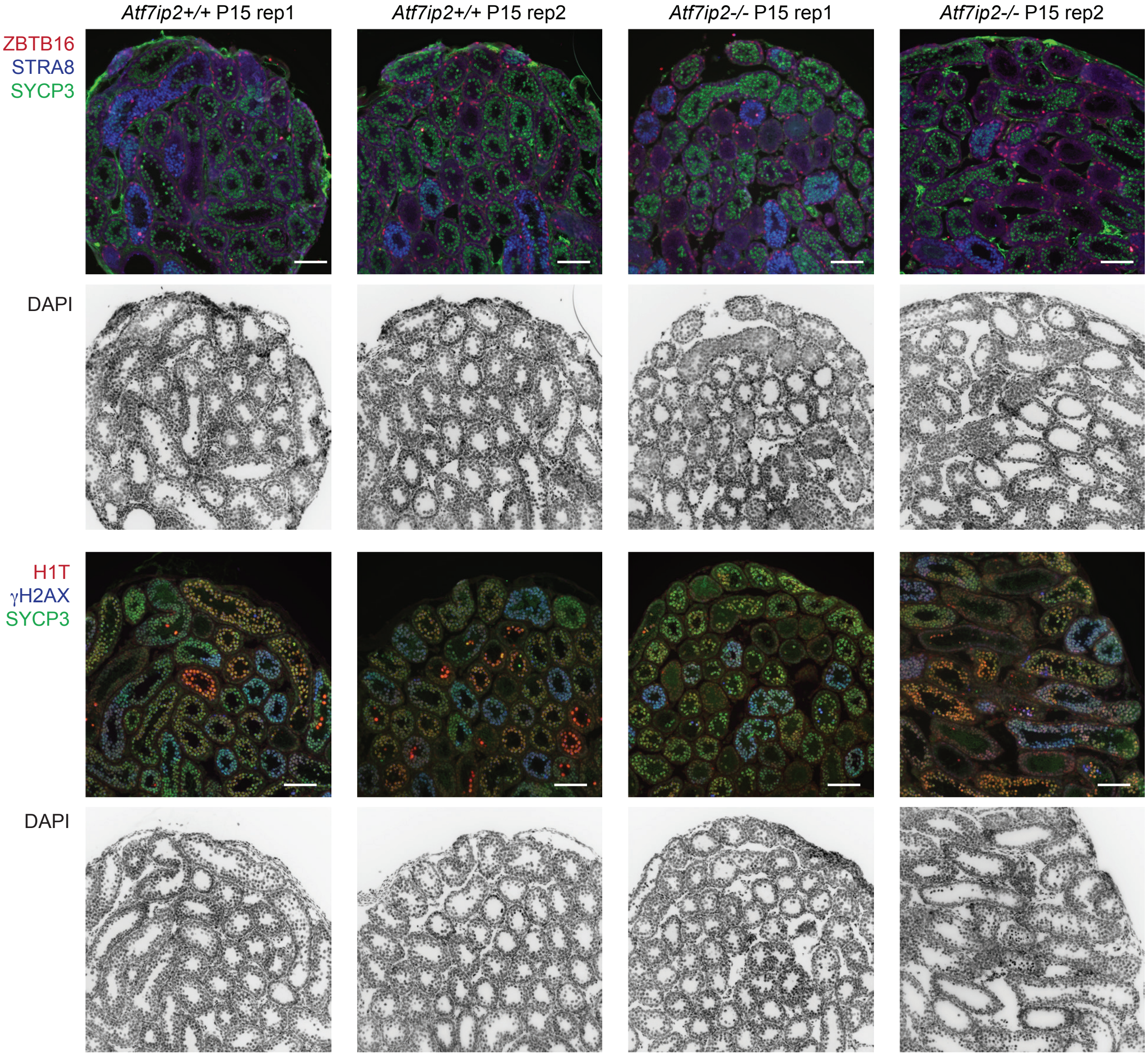


**Supplemental Figure S8. Testis sections from *Atf7ip2*$^{+/+}$ and *Atf7ip2*$^{-/-}$ mice at P15.**
Testis sections from *Atf7ip2*$^{+/+}$ and *Atf7ip2*$^{-/-}$ mice at P15 (used for scRNA-seq) stained with DAPI and antibodies raised against ZBTB16, STRA8, SYCP3, H1T, and γH2AX. Scale bars: 100 µm.

**A**

| | Platform: Chromium (10x Genomics) | |
|---|---|---|
| Genotype: | *Atf7ip2+/+* | *Atf7ip2-/-* |
| Sampling stage: | P15 | |
| Cell type: | Whole testicular cells | |
| Number of used indivisuals: | 2 | 3 |
| Total number of genes detected: | 26,168 | 26,415 |
| Sequencing Saturation: | 51.5% | 49.5% |
| Mean reads / cell: | 58,206 | 63,564 |
| Median genes / cell: | 4,174 | 4,566 |
| Median UMI counts / cell: | 14,666 | 16,156 |
| Total number of cells: | 9,113 | 9,015 |
| Number of analyzed testicular germ cells: | 959 (10.5%) | 2,246 (24.9%) |

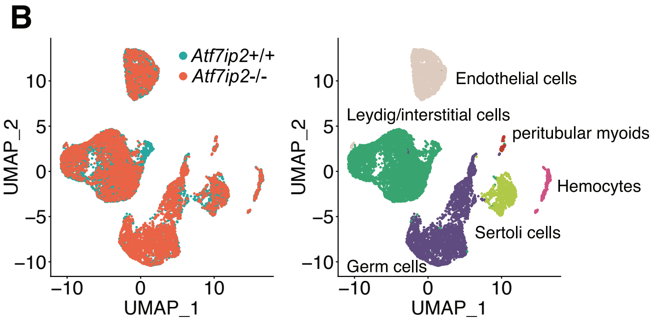


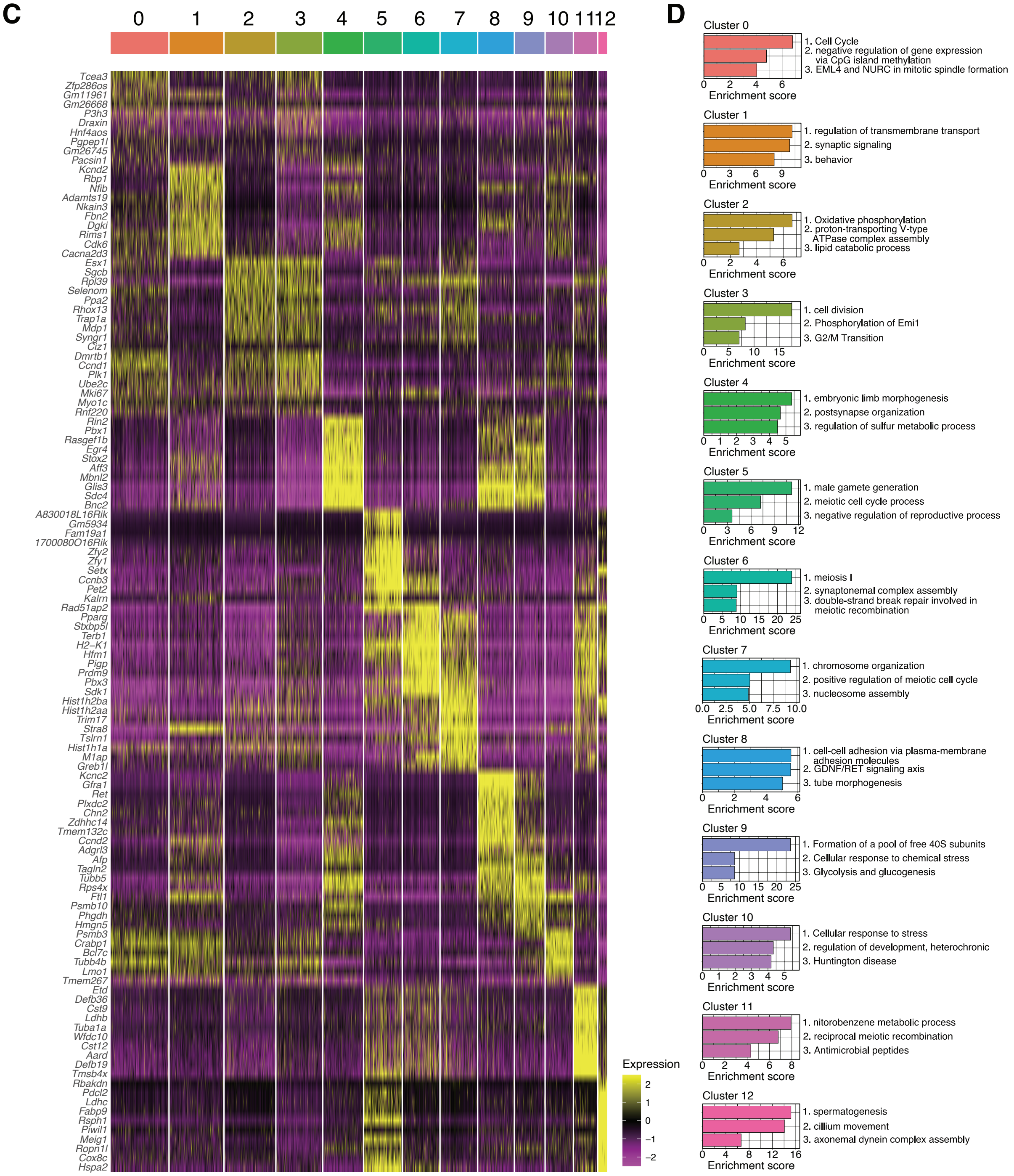

**Supplemental Figure S9. Clustering analysis of scRNA-seq data of *Atf7ip2*$^{+/+}$ and *Atf7ip2*$^{-/-}$ spermatogenic germ cells**

**(A)** Table of 10X Genomics Chromium metrics for scRNA-seq analyses of *Atf7ip2*$^{+/+}$ and *Atf7ip2*$^{-/-}$ testicular germ cells. For the *Atf7ip2*$^{+/+}$ sample, two *Atf7ip2*$^{+/+}$ testes from two individuals (P15) were merged. For the *Atf7ip2*$^{-/-}$ sample, three testes from three *Atf7ip2*$^{-/-}$ individuals (P15) were merged. We show the total numbers of testicular germ cells subjected to scRNA-seq analyses and separated from other testicular somatic cells. We also show percentages of extracted testicular germ cells per total single cells subjected to scRNA-seq analyses.

**(B)** UMAP representation of scRNA-seq transcriptome profiles for whole testicular cells from *Atf7ip2*$^{+/+}$ and *Atf7ip2*$^{-/-}$ testes.

**(C)** Hierarchically clustered heatmap for the top 10 representative genes for each cluster. Relative expression levels are indicated with the color code: yellow, up two positive standard deviations; black, mean; purple, up to two negative standard deviations.

**(D)** Gene enrichment analysis for genes in the cell clusters.

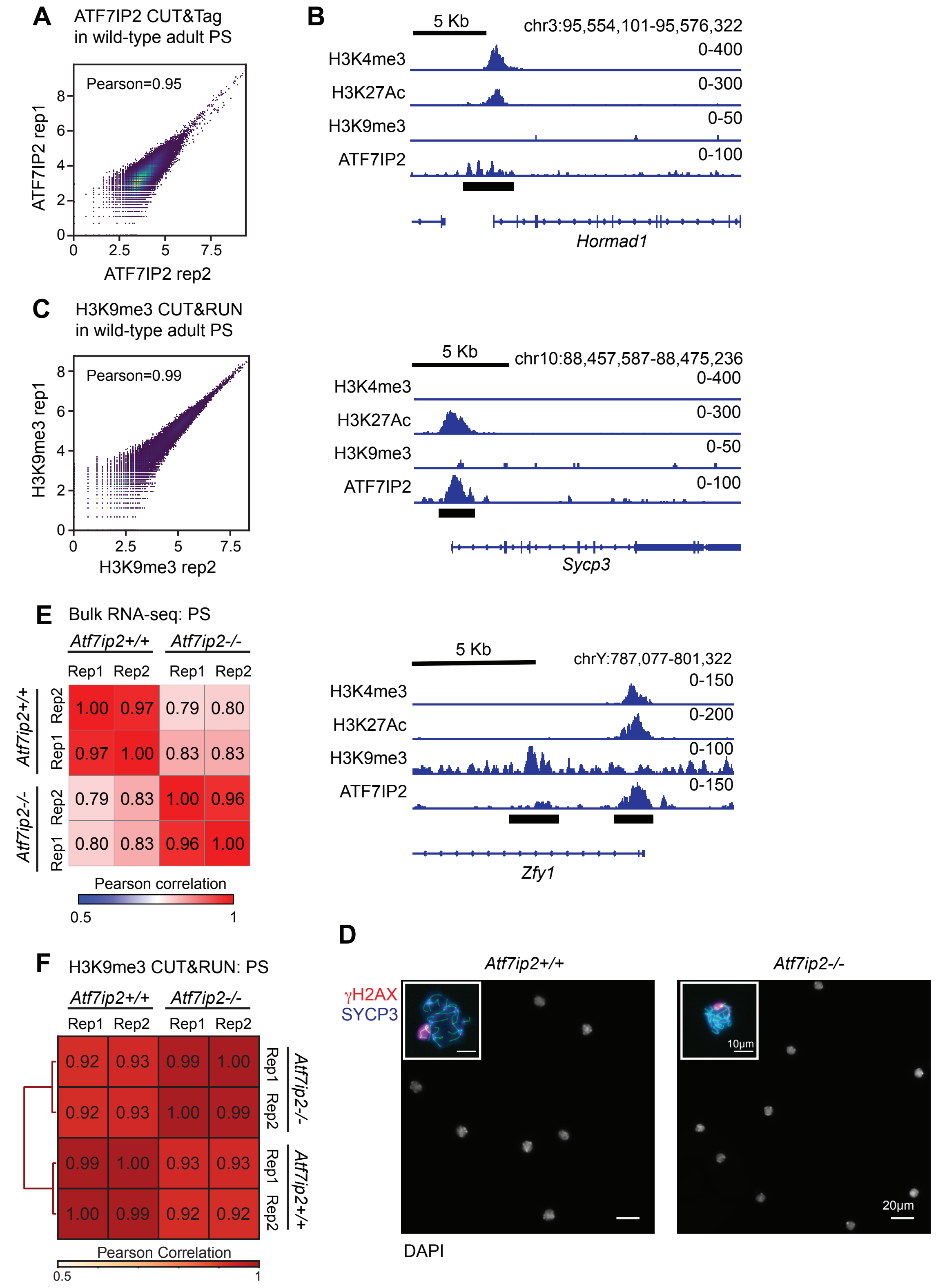
A
ATF7IP2 CUT&Tag
in wild-type adult PS
Pearson=0.95
ATF7IP2 rep1
ATF7IP2 rep2
0
2.5
5
7.5
2
4
6
8
B
5 Kb
chr3:95,554,101-95,576,322
H3K4me3
0-400
H3K27Ac
0-300
H3K9me3
0-50
ATF7IP2
0-100
Hormad1
C
H3K9me3 CUT&RUN
in wild-type adult PS
Pearson=0.99
H3K9me3 rep1
H3K9me3 rep2
5 Kb
chr10:88,457,587-88,475,236
H3K4me3
0-400
H3K27Ac
0-300
H3K9me3
0-50
ATF7IP2
0-100
Sycp3
E
Bulk RNA-seq: PS
Atf7ip2+/+
Atf7ip2-/-
Rep1 Rep2 Rep1 Rep2
Rep2 1.00 0.97 0.79 0.80
Rep1 0.97 1.00 0.83 0.83
Rep2 0.79 0.83 1.00 0.96
Rep1 0.80 0.83 0.96 1.00
Pearson correlation
0.5
1
5 Kb
chrY:787,077-801,322
H3K4me3
0-150
H3K27Ac
0-200
H3K9me3
0-100
ATF7IP2
0-150
Zfy1
D
Atf7ip2+/+
Atf7ip2-/-
γH2AX
SYCP3
10µm
20µm
DAPI
F
H3K9me3 CUT&RUN: PS
Atf7ip2+/+
Atf7ip2-/-
Rep1 Rep2 Rep1 Rep2
0.92 0.93 0.99 1.00 Rep1
0.92 0.93 1.00 0.99 Rep2
0.99 1.00 0.93 0.93 Rep1
1.00 0.99 0.92 0.92 Rep2
Pearson Correlation
0.5
1

**Supplemental Figure S10. ATF7IP2 CUT&Tag, H3K9me3 CUT&RUN, and bulk RNA-seq analyses in pachytene spermatocytes.**

**(A)** Scatter plots showing Pearson correlations between biological replicates for ATF7IP2 CUT&Tag data from wild-type adult pachytene spermatocytes.

**(B)** Representative track views of ATF7IP2 peaks at autosomal *Hormad1* and *Sycp3* gene loci, and the Y-linked *Zfy1* gene locus. Solid bars: locations of detected peaks.

**(C)** Scatter plots showing Pearson correlations between biological replicates for H3K9me3 CUT&RUN data from wild-type adult pachytene spermatocytes.

**(D)** Representative photos of isolated pachytene spermatocytes stained with DAPI; scale bars: 20 µm. Insets: Representative photos of isolated pachytene spermatocytes stained with antibodies raised against γH2AX and SYCP3; scale bars: 10 µm.

**(E)** Heatmap of Pearson correlations demonstrating the reproducibility of RNA-seq biological replicates.

**(F)** Heatmap of Pearson correlations demonstrating the reproducibility of H3K9me3 CUT&RUN biological replicates.

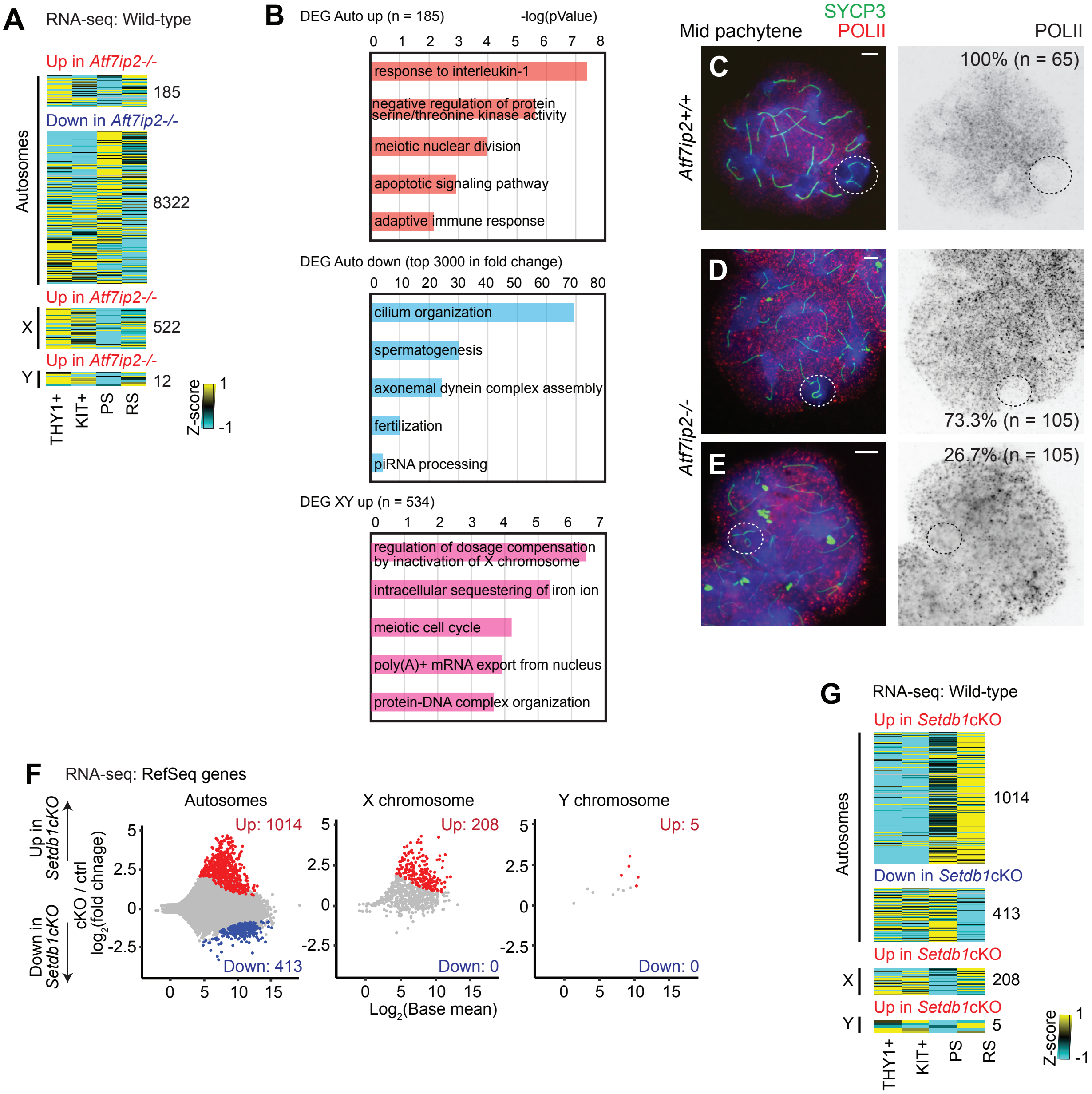


**Supplemental Figure S11. The roles of ATF7IP2 and SETDB1 in meiotic gene regulation**

**(A)** Heatmaps showing normalized expression of DEGs in *Atf7ip2*$^{+/+}$ and *Atf7ip2*$^{-/-}$ wild-type pachytene spermatocytes: THY1$^{+}$, undifferentiated spermatogonia; KIT$^{+}$, differentiating spermatogonia; PS, pachytene spermatocytes; RS, round spermatids.

**(B)** The top five Gene Ontology enrichment terms for autosomal up- and downregulated genes and XY-linked upregulated genes from differential expression analyses of *Atf7ip2*$^{-/-}$ and *Atf7ip2*$^{+/+}$ pachytene spermatocyte RNA-seq.

**(C, D, E)** *Atf7ip2*$^{+/+}$ and *Atf7ip2*$^{-/-}$ spermatocyte chromosome spreads stained with antibodies raised against SYCP3 (a marker of chromosome axes) and RNA polymerase II (POLII). Three independent littermate pairs were examined. Numbers of analyzed nuclei are shown. Scale bars: 10 µm. Dashed circles indicate the sex chromosomes.

**(F)** Comparison of *Setdb1* control (*Setdb1* ctrl) and *Setdb1* conditional knockout (*Setdb1*-cKO) pachytene spermatocyte transcriptomes. Autosomal, X, and Y genes were analyzed separately. Two independent biological replicates were examined. All genes with adjusted p-values (Benjamini-Hochberg method) are plotted. Differentially expressed genes (DEGs: fold change ≥ 1.5, adjusted p-value ≤ 0.05) are colored (red: upregulated in *Setdb1*-cKO; blue: downregulated in *Setdb1*-cKO), and numbers are shown.

**(G)** Heatmaps showing normalized expression of DEGs in *Setdb1* ctrl and *Setdb1*-cKO wild-type pachytene spermatocytes.

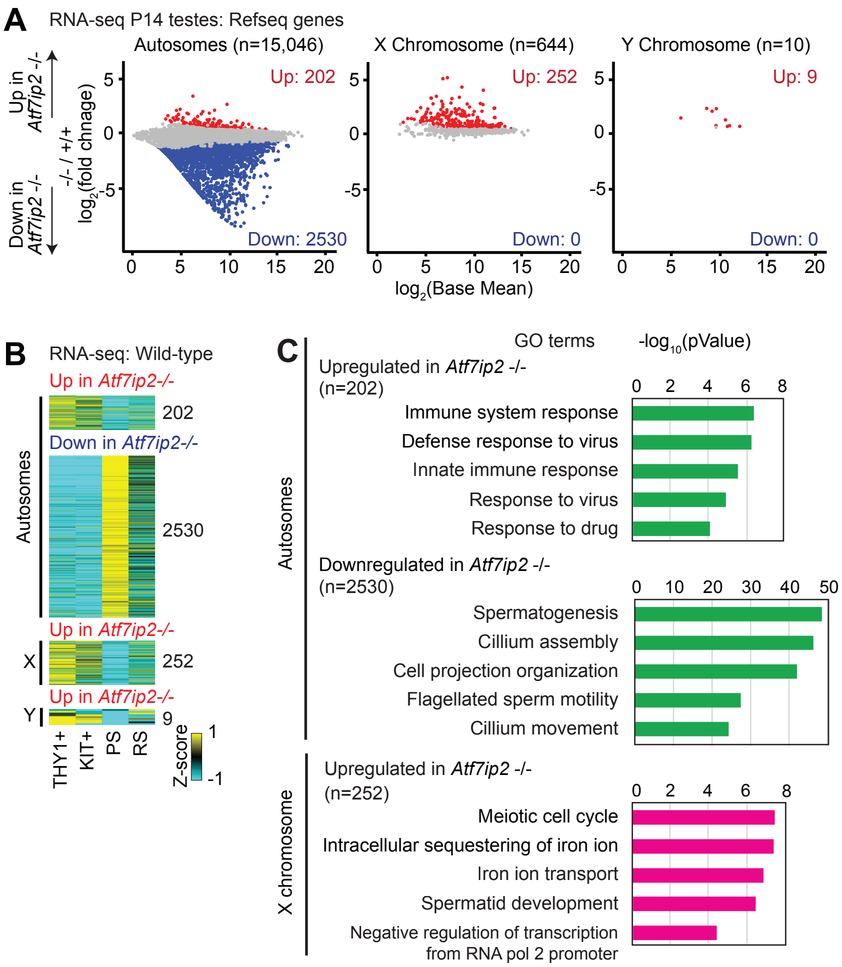


**Supplemental Figure S12. RNA-seq analysis of *Atf7ip2*$^{+/+}$ and *Atf7ip2*$^{-/-}$ testes at P14.**

**(A)** Comparison of transcriptomes in P14 *Atf7ip2*$^{+/+}$ and *Atf7ip2*$^{-/-}$ testes. Autosomal, X, and Y genes were separately analyzed. Two independent biological replicates were examined. All genes with adjusted p-values (Benjamini-Hochsberg method) are plotted. Differentially expressed genes (DEGs: fold change ≥ 1.5, adjusted p-value ≤ 0.05) are colored (red: upregulated in *Atf7ip2*$^{-/-}$ testes; blue: downregulated in *Atf7ip2*$^{-/-}$ testes), and numbers are shown.

**(B)** Heatmaps showing normalized expression of DEGs between *Atf7ip2*$^{+/+}$ and *Atf7ip2*$^{-/-}$ testes in wild-type spermatogenesis: THY1$^{+}$, undifferentiated spermatogonia; KIT$^{+}$, differentiating spermatogonia; PS, pachytene spermatocytes; RS, round spermatids.

**(C)** The top five Gene Ontology enrichment terms for autosomal up- and downregulated genes and X-linked upregulated genes from differential expression analyses of *Atf7ip2*$^{-/-}$ and *Atf7ip2*$^{+/+}$ P14 testis RNA-seq.

**Table S1 ScRNA-seq analysis**
Top 100 differentially expressed genes for each cluster and results of gene enrich analysis for each cluster

**Table S2 ATF7IP2 binding target genes determined by the ATF7IP2 CUT&Tag analysis.**
Lists of autosomal ATF7IP2 binding target genes with which ATF7IP2 binds on promoters.
Lists of XY-linked ATF7IP2 binding target genes with which ATF7IP2 binds on promoters.
List of XY-linked ATF7IP2 binding target genes with which ATF7IP2 binds entire gene bodies.

**Table S2 Differentially expressed genes between *Atf7ip2*$^{+/+}$ and *Atf7ip2*$^{-/-}$ pachytene spermatocytes**
List of differentially expressed genes (DEG) determined after ERCC normalization.
